# Cross-Species Multi-Omics Profiling Identifies Conserved Activated Valvular Interstitial Cell Population Driving Myxomatous Mitral Valve Degeneration

**DOI:** 10.64898/2026.03.23.713796

**Authors:** Fu Gao, Ian Mason, Mingze Dong, Yao Lu, Di Zhang, Xing Lou, Irbaz Hameed, Mingyu Yang, Mei Zhong, Markus Krane, Giovanni Ferrari, George Tellides, Yang Liu, Rong Fan, Arnar Geirsson

## Abstract

**Background:** Primary mitral regurgitation resulting from mitral valve prolapse can lead to life-threatening complications, including arrhythmias, heart failure, and sudden cardiac death. Mitral valve prolapse is classically associated with myxomatous mitral valve degeneration, characterized by leaflet thickening, extracellular matrix disorganization, and progressive structural remodeling. Valvular interstitial cells, the predominant stromal population within the valve, maintain extracellular matrix homeostasis; however, their molecular heterogeneity, and state-specific contributions to disease pathogenesis remain incompletely defined.

**Methods:** Using a fibrillin-1 deficient mouse model and human tissue specimens we integrated single-cell RNA sequencing with spatial transcriptomic profiling to construct a comprehensive atlas of cellular composition and extracellular matrix organization across normal mitral valves, sporadic mitral valve prolapse, and Marfan syndrome-associated mitral valve prolapse.

**Results:** Analyses revealed spatially organized cellular niches and substantial heterogeneity within the valvular interstitial cell population. Across murine and human datasets, we identified a conserved activated valvular interstitial cell population enriched for profibrotic extracellular matrix remodeling programs and preferentially localized to mechanically vulnerable leaflet tip regions. This population exhibited coordinated upregulation of collagen- and matrix-associated genes, metabolic signatures consistent with enhanced mitochondrial activity, and transcriptional features suggesting fibro-inflammatory signaling.

**Conclusions:** We identified a transcriptionally and spatially distinct activated valvular interstitial cell state conserved across species and disease etiologies that is strongly implicated in fibrotic remodeling during myxomatous mitral valve degeneration and provides a candidate therapeutic target.

## Introduction

Mitral valve disease imposes a substantial global health burden and can lead to progressive heart failure, arrhythmia, infective endocarditis, and sudden cardiac death (1). Sporadic mitral valve prolapse (MVP), affecting approximately 1-2% of the population, is the leading cause of primary mitral regurgitation in adults and is most commonly associated with myxomatous mitral valve degeneration (MMVD) (2). Among syndromic forms, Marfan syndrome-associated MVP is the most prevalent, characterized by more pronounces MMVD involving both mitral valve leaflets, and carries a substantially increased likelihood of requiring surgical intervention compared with non-syndromic MVP (3). Although surgical management of mitral regurgitation has advanced considerably, favoring valve repair over replacement and incorporating transcatheter techniques, disease-modifying medical therapies remain unavailable. This therapeutic gap largely reflects an incomplete understanding of the cellular and molecular mechanisms that drive progressive leaflet remodeling.

Histopathologically, MMVD is characterized by leaflet thickening and elongation, disorganization of the extracellular matrix (ECM) architecture, and excessive accumulation of collagen- and glycosaminoglycans- (GAGs) rich matrix (4). Although the disease has historically been described as “myxomatous”, the contribution of fibrosis-associated ECM remodeling to disease progression remains insufficiently defined. This gap in knowledge partly reflects the technical difficulty of systematically profiling the mitral leaflet, whose thin, elongated structure complicates whole-leaflet spatial analysis (5). Consequently, prior investigations have predominantly examined discrete or regionally restricted valve regions. As a result, comprehensive histologic characterization of the entire human mitral leaflet remains incomplete, and a spatially resolved, whole-organ molecular atlas of normal and diseased human mitral leaflets has not been established.

Valvular interstitial cells (VICs), mesenchyme-derived fibroblast-like stromal cells, constitute the predominant resident cellular population within the mitral valve and are central regulators of ECM homeostasis and mechanical integrity (6–8). Although fibroblast heterogeneity has been extensively described in multiple organs, the diversity and functional specialization of mitral VICs remains incompletely defined. Traditionally viewed as quiescent in the healthy valve, VICs are now recognized to exhibit dynamic transcriptional and metabolic states even under physiological conditions (7,9). Under pathological stress, VICs can adopt activated matrix-remodeling phenotype characterized by increased collagen synthesis, induction of matrix metalloproteinases, and upregulation of inflammatory mediators and chemokines (10–12). Whether specific VIC states spatially organize and drive the fibrotic remodeling observed in MMVD remains unclear.

In this study, we integrated single-cell RNA sequencing (scRNA-seq), spatial transcriptomics, and ECM mapping to construct a spatially resolved cellular and molecular atlas of the mitral valve. Using a murine model together with human specimens spanning normal, sporadic MVP, and Marfan-associated MVP, we identify multiple transcriptionally and spatially distinct VIC subtypes. Notably, under diseased conditions, we defined a conserved activated VIC population enriched for profibrotic ECM-remodeling program while showing limited induction of canonical contractile markers traditionally associated with myofibroblast differentiation (13,14). These findings refine the cellular framework of MMVD pathogenesis by identifying a spatially restricted activated VIC state as a central effector of fibrotic remodeling and a candidate target for disease-modifying therapy.

## Methods

### Ethics

Animal research was approved by the Institutional Animal Care and Use Committee of Yale University. All mice were housed in standardized conditions within Yale University’s Animal Facility. All procedures were conducted in compliance with US legislative requirements. The study adhered to the US code of conduct for responsible human tissue use. The collection of the human tissue was approved by the Institutional Review Board of Yale University and the New England Organ Bank. All patients provided informed consent.

### Mice

*Fbn1^C1039G/+^* mice (C1039G) were a gift from Harry C. Dietz (available from Jackson Laboratory as stock no. 012885) (15). Mice were euthanized at 12 weeks old for analysis. Wild-type control hearts were harvested from gender-matched littermates in each group. After euthanasia, the thoracic cavity was extensively opened, and the hearts were perfused with saline. Subsequently, the mitral valve was isolated through a longitudinal incision and excised for subsequent experiments.

### Echocardiography

Mitral valve function was assessed using high-resolution echocardiography on animals anesthetized with a light isoflurane dose. A 40 MHz linear array transducer (Vevo 2100, VisualSonics) was employed to detect mitral regurgitation.

### Gross morphology examination

Mouse hearts were excised and fixed in 4% paraformaldehyde (PFA) at 4°C overnight. Following fixation, the mitral valves were exposed for examination. In situ images of the mitral valves were captured using an SZX16 stereoscopic microscope equipped with an Olympus camera. Human diseased mitral valve tissues were procured from patients undergoing repair surgery, while normal specimens were sourced from non-transplanted donor hearts. The control samples were selected based on the absence of mitral valve disease and overall health status to serve as a baseline for comparison with diseased samples. Photographs with a scale reference were taken immediately following acquisition.

### Histomorphometry

Tissues were fixed in 4% PFA overnight, paraffin-embedded, and sectioned at 7 μm thickness. Yale’s Research Histology Lab stained the sections using Hematoxylin and Eosin (H&E), Masson’s Trichrome (Trichrome), Von Koss, Alizarin red, and Movat’s Pentachrome (Movat) staining using standard techniques. Morphometric analysis was conducted with ImageJ.

### Confocal imaging

Tissues were embedded in OCT and sectioned at 7 μm thickness. After washing three times with Tris-buffered saline (TBS), tissue sections were incubated with primary antibodies diluted in blocking solution (10% BSA and horse serum in TBS) overnight at 4°C in a humidified enclosure. Sections were washed three times with TBS, incubated with corresponding Alexa Fluor 488-, Alexa Fluor 594-, or Alexa Fluor 647-conjugated secondary antibodies in blocking solution for 1 hour at room temperature, followed by another series of three TBS washes, and mounted on slides with ProLong Gold mounting reagent with DAPI (ThermoFisher Scientific, P36935). The following antibodies were used: anti-Tnfrsf12a (Invitrogen, 14-9018-82), anti-CD68 (Abcam, ab955), anti-TGF-β1 (Abcam, Ab92486), anti-CD45 (BD, 550539), anti-Collagen III (Abcam, ab7778), anti-CD68 (Bio-rad, MCA1957), anti-CD45 (R&D, AF114), anti-Collagen I (Abcam, ab34710), anti-RSPO3 (Proteintech, 17193-1-AP), anti-p-SMAD2 (Cell Signaling, 3108), anti-Elastin (Abcam, ab21610), anti-CD31 (R&D, AF3628), anti-CHAD (Invitrogen, PA5-53761), anti-Ifit3 (Proteintech, 15201-1-AP), anti-CCR2 (Abcam, ab216863), anti-LAP (R&D, MAB7666). All immunofluorescence images were taken with a Leica SP8 confocal microscope. ImageJ software was used to quantify immunofluorescence signals by calculating the mean fluorescence intensity, defined as the sum of pixel values within each valve section normalized to the measured area and expressed as arbitrary units (AU) or number of positive stained cells for specific antibodies.

### Single-cell RNA sequencing

Mitral valves were procured, rinsed in cold PBS, and sliced into small fragments. The minced tissue was incubated in DMEM with 1.5 mg/ml collagenase A, and 0.5 mg/ml elastase for 60 min at 37 °C. The digested solution was passed through a 70 μm filter and incubated with cell-impermeant viability dye (ThermoFisher Scientific, 65-0865-14) for 20 min, washed, resuspended in 0.4% BSA/PBS for sorting using a LSR II (BD Biosciences) as described (16). Single-cell suspensions were processed using the Chromium Controller (10x Genomics) according to the manufacturer’s instructions. Following cDNA amplification and library construction, sequencing was performed on the Illumina HiSeq 4000 platform at the Yale Center for Genome Analysis. Post-sequencing, the data for each specimen was aligned to the reference genomes of human or mouse as supplied by 10X Genomics as described (16) and employing the CellRanger suite following the default parameters.

### scRNA sequencing and raw data processing

scRNA-seq data was further processed in R using Seurat (4.3.0). Cells were filtered to include those with mitochondrial reads less than 10% and nCount_RNA between 500 and 15,000 (17,18). The gene expression of the remaining cells was normalized by the method “LogNormalize”. Highly variable genes were selected using standard variation and were used in the downstream analyses. Principal component analysis (PCA) was conducted based on highly variable genes for dimensionality reduction and 50 significant principal components were chosen for batch effect correction using Harmony. Clustering in Harmony was performed using graph-based clustering approach with an appropriate resolution for each data. The Louvain algorithm was used to group cells into different cluster. Uniform Manifold Approximation and Projection (UMAP) were applied for the two-dimensional visualization of the clustering all cell types and specific cell subtypes respectively. Differentially expressed genes were identified by Wilcoxon test. Genes with log2FC (fold change, FC) >0.25 and adjusted p-value <0.05 were considered as significant differentially expressed genes.

### scRNA-seq data analysis

To evaluate the ligand-receptor interactions among the identified cell types, we retrieved the gene expression matrix from the RDS files, annotated with cell type information. Subsequently, we applied CellChat to infer the ligand-receptor interactions as previously described (19). Single-cell metabolic data were analyzed using the scMetabolism package on identified cell types to quantify the metabolic activity at the single cell resolution. Senescence signature scores were computed using the SenMayo gene set (20) (Supplemental Tabel 1) and the AddModuleScore function implemented in Seurat.

### Spatial transcriptome profiling of mouse and human mitral valve

Two types of chips were used in this study, the 50x50 10um chip was used for mouse mitral valve spatial transcriptome profiling and 100x100 20um chip was used for human mitral valve spatial transcriptome profiling (21,22).

### Analysis of spatial omics data

The library was built and sequenced by an illumina Novaseq 6000 sequencer.

For cDNAs originating from mRNAs, the raw FASTQ file, which includes the UMI, barcode A, and barcode B, was restructured into the format required by ST Pipeline version 1.7.2 using a custom Python script (23). Adhering to ST Pipeline’s suggested settings, the RNA expression matrices were clustered using Seurat (24). For the transcriptome data, normalization was performed using the SCTransform function in Seurat.

### Quantitative RT-PCR

Total RNA was isolated from mitral valve tissues using the RNeasy Mini kit (Qiagen, Inc. 74104). The quantification of target genes was performed by using the High-capacity cDNA Reverse Transcription kit (ThermoFisher Scientific, 4368814) and TaqMan Gene Expression Master Mix (ThermoFisher Scientific, 4369016) according to manufacturer’s instructions. The amplified genes and primer catalogs (ThermoFisher Scientific) were as follows: *COL1A1* (Hs00164004_m1), *COL1A2* (Hs01028956_m1), *COL3A1* (Hs00943809_m1), *COL5A1* (Hs00609133_m1), *COL6A1* (Hs01095585_m1), *COL10A1* (Hs00166657_m1), *COL11A1* (Hs01097664_m1), *COL11A2* (Hs00899176_m1), *COL13A1* (Hs01103890_m1), *COL14A1* (Hs00964045_m1), *COL15A1* (Hs00266332_m1), *COL21A1* (Hs00229402_m1), *MMP11* (Hs00968295_m1), *MMP16* (Hs00234676_m1), *ADAMTS14* (Hs01548440_m1), *ADAMTS16* (Hs00373526_m1), *ACAN* (Hs00153936_m1), *DCN* (Hs00754870_s1), *LUM* (Hs00929860_m1), *PRG4* (Hs00981633_m1), and *ACTB* (Hs99999903_m1). The amplification procedures were performed on Bio-Rad CFX94. The results were normalized to the expression of ACTB, and all data are expressed as 2^-ΔΔCt^.

### Micro-computed tomography

Mitral valve calcification was assessed using high-resolution micro-computed tomography (Scanco Medical AG, Brüttisellen, Switzerland) at the Yale microCT Imaging Facility. Human mitral valve tissues were scanned, and cross-sectional images were reconstructed using the manufacturer’s software.

### Statistics and visualization

Numerical data are depicted as point graphs showing individual observations, with lines indicating the average and standard error of the mean (SEM). Continuous variables across two cohorts were compared using the t-test, while comparisons among multiple groups utilized one-way ANOVA for the independent variable, with post-hoc Tukey’s tests applied when ANOVA indicated significant differences. P-values were bidirectional, and a threshold of P < 0.05 was set for statistical significance. All graphs and statistical evaluations were conducted using Prism version 10.0.0 (GraphPad Software). Symbols such as asterisks indicate statistical significance levels in group comparisons. Differential expression analysis was performed using Seurat’s FindMarkers function, employing the Wilcoxon test to identify differentially expressed genes between groups.

### Data and code availability

The datasets and custom code generated in this study are being prepared for public deposition and will be made available in a public repository prior to manuscript acceptance.

## Results

### Characterization of the *Fbn1^C1039G/+^* model of myxomatous mitral valve degeneration

The fibrillin-1 (Fbn1) deficient heterozygous *Fbn1^C1039G/+^* (C1039G) mouse is a well-established model of Marfan syndrome that develops aortic aneurysms, kyphosis, skeletal muscle myopathy while maintaining a near-normal lifespan. Importantly, this model also recapitulates key features of MVP (15,25,26). By 12 weeks of age, C1039G mice exhibited structural abnormalities closely resembling human MMVD, including leaflet enlargement, thickening, and architectural disorganization, most prominently at the leaflet tip (Figure 1A). These structural changes were accompanied by high rates of mitral regurgitation (Figure 1B). Immunofluorescence analysis revealed increased TGF-β1 and phosphorylated Smad2 (p-SMAD2) expression, together with reduced levels of latency-associated peptide (LAP) in C1039G valves compared with wild-type (WT, +/+) controls (Supplemental Figure 1), consistent with enhanced TGF-β pathway activation.

**Figure 1.**
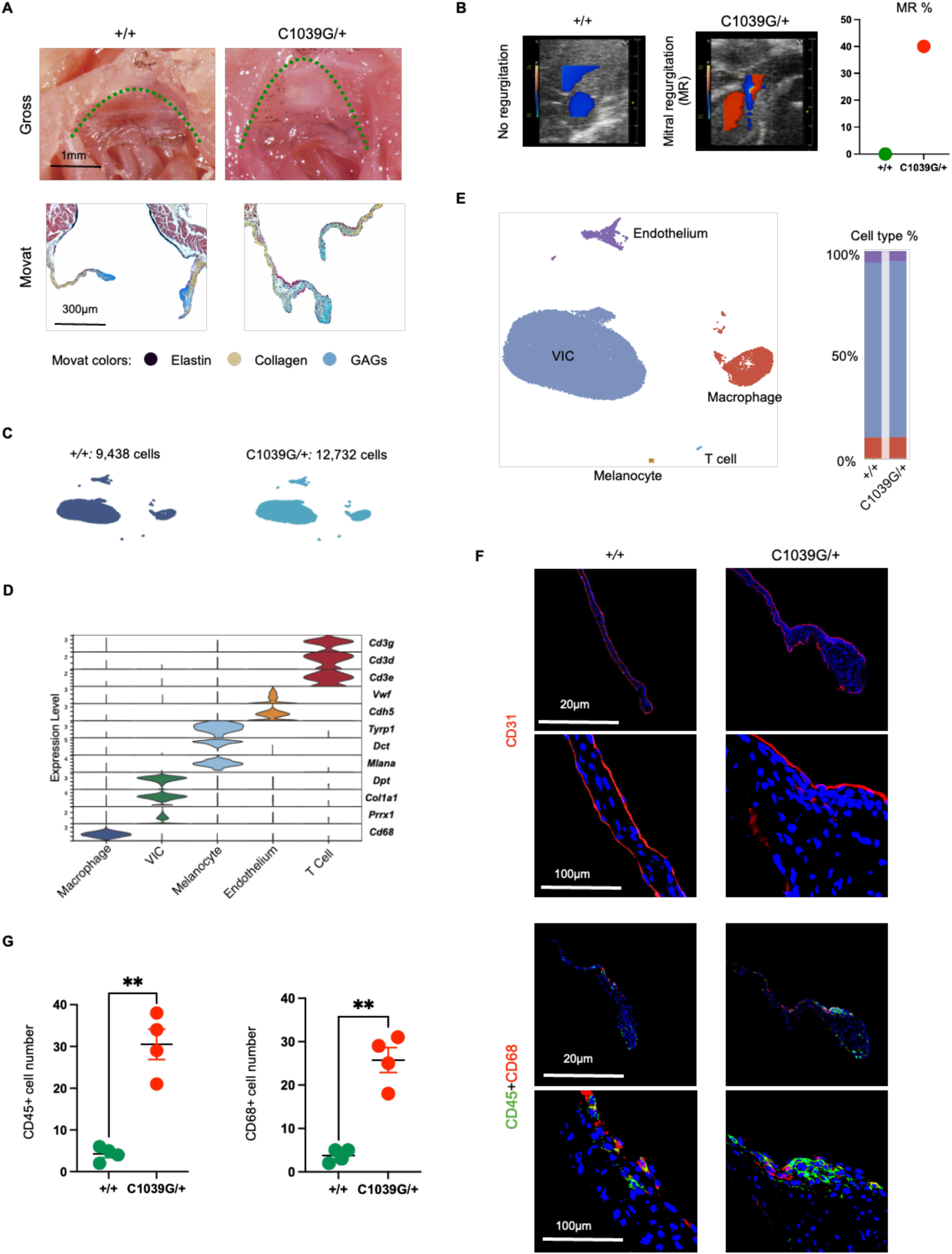
Structural and single-cell characterization of mitral valves in Fbn1-deficient mice. (**A**) Gross morphology of mitral valve leaflets (upper panels). Green dashed lines delineate leaflet boundaries. Representative Movat’s pentachrome staining (lower panels) showing extracellular matrix (ECM) composition and architectural organization. GAGs, glycosaminoglycans. (**B**) Representative echocardiographic images showing mitral regurgitation (MR) in *C1039G/+* mice compared with wild-type (+/+) controls, with corresponding quantification of MR percentage. (**C**) UMAP visualization of single-cell RNA-sequencing (scRNA-seq) data from +/+ and C1039G/+ mitral valves. (**D**) Violin plots showing expression of canonical marker genes across major annotated valve cell populations. (**E**) UMAP colored by cell type, with bar plots indicating the relative proportions of each cell population in +/+ and C1039G/+ valves. (**F-G**) Representative immunofluorescence images of mitral valve sections stained for endothelial cells (CD31), leukocytes (CD45), and macrophages (CD68), with quantitative analysis. Data are shown as individual data points with mean ± SEM. Statistical comparisons were performed using unpaired t test. **P<0.01.

We next performed scRNA-seq on mitral valves from C1039G and WT mice. Because of the small size and low cellularity of the murine mitral leaflet, valves from 20 mice per group were pooled to obtain sufficient cells input, yielding 22,170 high-quality cells after quality control (Figure 1C). Based on established lineage-specific marker genes, five major cell types were identified: VICs, macrophages, T cells, endothelial cells, and melanocytes (Figure 1D-E) (17,18). Immunofluorescence staining demonstrated spatial distribution of these cell types within the leaflet. A CD31^+^ endothelial monolayer lined the leaflet surface, whereas CD68^+^ macrophages localized predominantly to the atrial side of the mid-leaflet and to the leaflet tip. The number of macrophage was increased in C1039G valves compared with WT controls (Figure 1F-G). A small melanocytes population was also detected by scRNA-Seq in murine mitral valves, although its functional contribution to valve biology remains undefined.

### Murine mitral valve VICs comprise distinct functional subtypes with disease-associated shifts in abundance and spatial distribution

Unsupervised clustering of VICs from WT and C1039G mitral valves revealed marked transcriptional heterogeneity, with five distinct subsets (mVIC1-mVIC5) (Figure 2A-B). To define their functional characteristics, we examined genes associated with ECM synthesis and inflammatory signaling. In WT valves, each VIC subset displayed a distinct transcriptional profile, reflected by differential expression of structural ECM components, including collagens (*Col1a1, Col3a1, Col8a1*), elastin (*Eln*), GAG biosynthetic enzymes (*Has2, Ugdh, Ugp2*), inflammatory mediators (*Cxcl2, Ccl2*), and the chemokine receptor *Ccr2* (Figure 2C).

**Figure 2.**
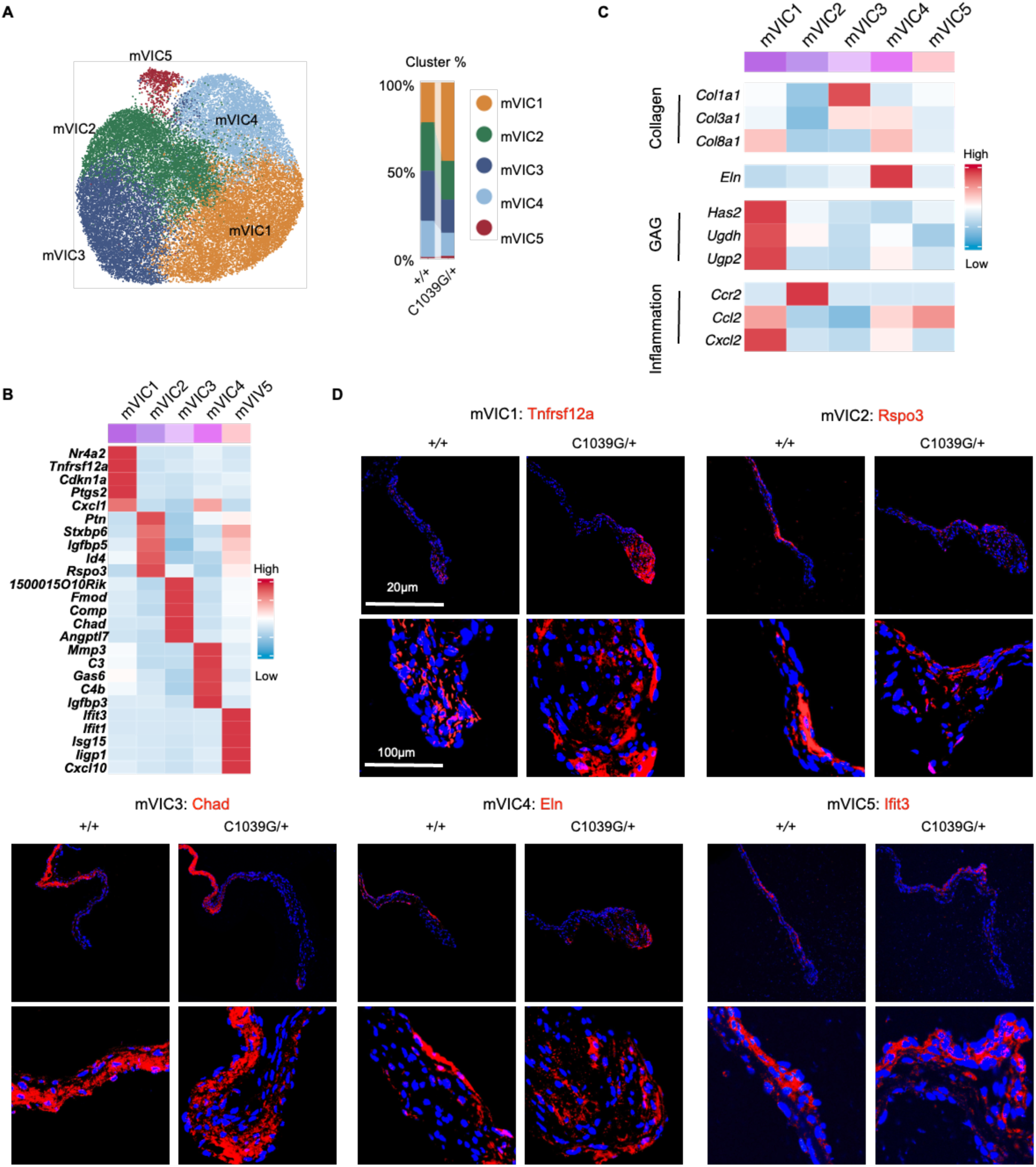
Transcriptional and spatial characterization of VIC subsets in Fbn1-deficient mice. (**A**) UMAP projection showing unsupervised clustering of valvular interstitial cells (VICs). Bar plots indicate the relative proportion of each VIC cluster in +/+ and C1039G/+ valves. (**B**) Heatmap showing the top five differentially expressed genes (DEGs) in each VIC cluster in +/+ and C1039G/+ valves. (**C**) Heatmap of representative genes across VIC clusters in WT valves only, highlighting extracellular matrix (ECM) biosynthetic programs and proinflammatory transcriptional signatures. (**D**) Representative immunofluorescence images demonstrating the spatial distribution of distinct VIC subsets within mitral valve leaflets.

mVIC1 exhibited concurrent enrichment of GAG-related and inflammatory transcripts and was significantly expanded in C1039G valves compared with WT controls (Figure 2A and 2C). In contrast, mVIC3 and mVIC4 were primarily enriched for ECM programs, with mVIC3 characterized by elevated collagen expression and mVIC4 preferentially expressing elastin-associated transcripts. mVIC2 and mVIC5 showed relatively higher expression of immune-associated genes (Figure 2C). Together, these findings support the presence of functionally specialized VIC subtypes that likely contribute to regional ECM composition and immune modulation within the WT valve.

To determine whether these transcriptionally defined VIC clusters exhibited spatial organization across the leaflet, we performed immunofluorescence staining using representative markers selected from cluster-enriched genes (Figure 2B). The mVIC1 subset (Tnfrsf12a⁺) was predominantly localized to the leaflet tips and was markedly increased in C1039G valves compared with WT controls (Figure 2D).

Consistent with elevated *Ccl2* expression in mVIC1 from C1039G valves (Figure 2C), C1039G mice exhibited increased accumulation of CD45⁺CCR2⁺ immune cells (Supplemental Figure 2) (12). The mVIC2 (Rspo3⁺) exhibited a distinct inflammatory transcriptional profile and was primarily distributed within the mid-leaflet region in WT valves, while in C1039G, this population was increased and displayed spatial overlap with macrophages-enriched regions (Figure 1F). mVIC3 (Chad⁺), enriched for collagen transcripts, was localized predominantly at the leaflet base, consistent with collagen-dense regions observed by Movat’s pentachrome staining (Figure 1A). The elastin-enriched subset mVIC4 (Eln⁺) exhibited a broad distribution extending from the leaflet base toward the proximal tip, with higher density along the atrial side, consistent with elastin-rich zones observed by Movat’s staining (Figure 1A). Finally, mVIC5 (Ifit3⁺), characterized by inflammatory-associated gene expression, localized predominantly to the mid-leaflet and overlapped with leukocyte-rich regions (Figure 1F).

### Activated murine VIC subset drives MMVD remodeling

Compared with WT valves, C1039G valves exhibited the most pronounced structural alterations at the leaflet tip, rather than at the leaflet base or mid-leaflet (Figures 1A and 3A), paralleling the regional pattern observed in human clinical MMVD (5). Movat’s pentachrome staining further showed that the leaflet tip is enriched in GAGs under baseline conditions (Figure 1A), whereas in C1039G valves this region displayed marked ECM disorganization and collagen accumulation (Figures 1A and 3A). Because mVIC1 preferentially localized to the leaflet tip (Figure 2D), was strongly associated with GAG biosynthetic programs (Figure 2C), and was expanded in C1039G valves (Figure 2A), we next examined this population in greater detail in the diseased state.

**Figure 3.**
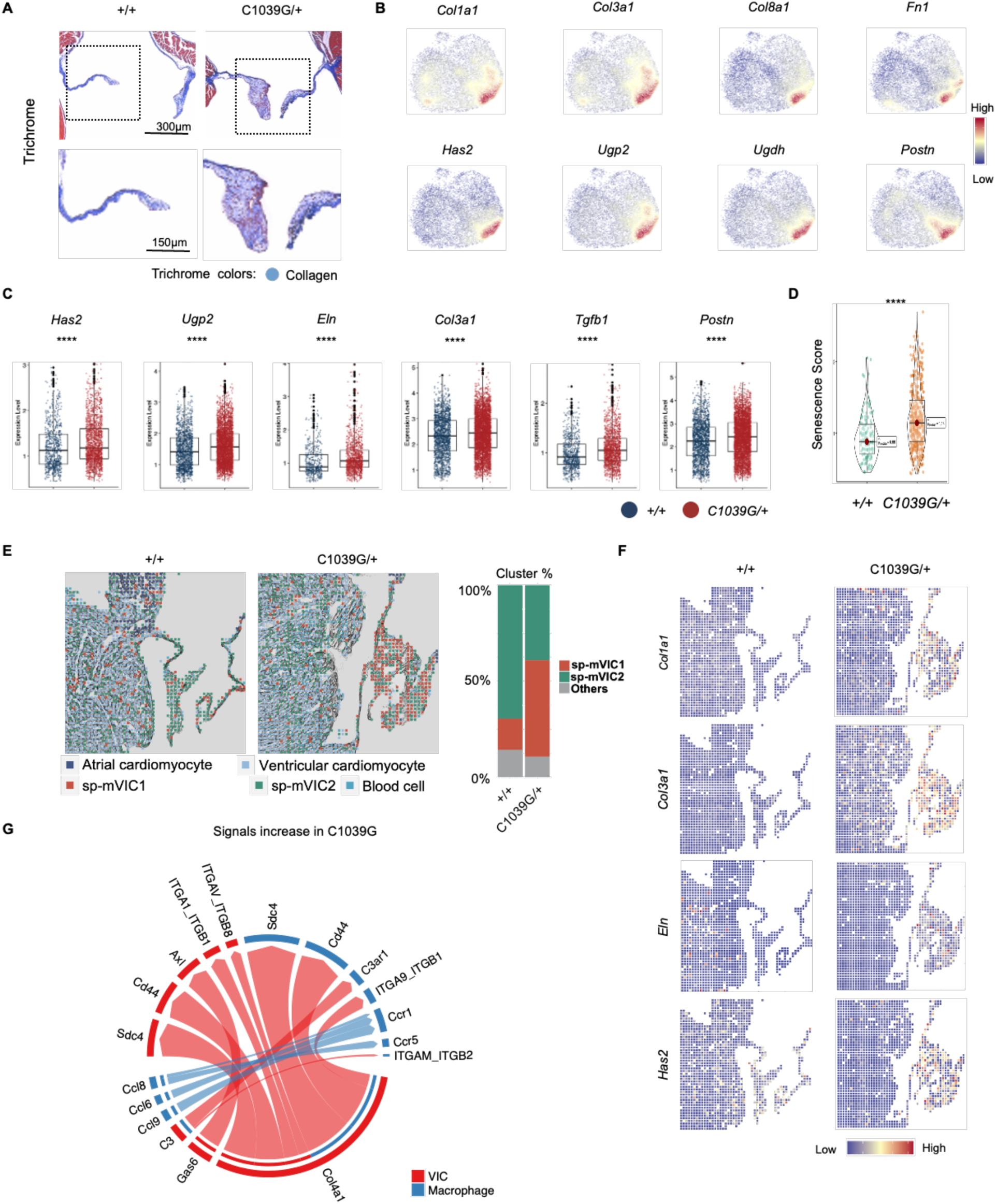
Transcriptional and spatial characterization of VIC subsets in Fbn1- deficient mice. (**A**) Representative Masson’s Trichrome (Trichrome) staining showing marked morphological changes at the leaflet tip in C1039G/+ mice compared with +/+ controls. (**B**) UMAP showing enrichment of representative ECM and profibrotic genes within the mVIC1 cluster. (**C**) Boxplots quantifying expression of selected genes in mVIC1 comparing +/+ and C1039G/+ valves. Data are shown as individual data points with mean ± SEM. ****P < 0.0001, by Wilcoxon test. (**D**) Violin plots comparing senescence-associated gene set scores in mVIC1 from +/+ and C1039G/+ mice. Data are presented as individual data points with mean ± SEM. ****P < 0.0001, by Wilcoxon test. (**E**) Spatial transcriptomic mapping of unsupervised clusters in mitral valves from +/+ and C1039G/+ mice. Bar plots shows the relative proportions of spatial clusters within the mitral valve leaflets. (**F**) Spatial transcriptomic feature maps demonstrating expression of major ECM-related genes in +/+ and C1039G/+ valves. (**G**) Representative ligand-receptor interaction networks upregulated in C1039G/+ compared with +/+ mitral valves. Edge thickness represents inferred interaction strength.

In WT valves, distinct VIC subsets displayed a balanced and homeostatic distribution of ECM and inflammatory programs, with different subsets preferentially contributing specific matrix components and signaling mediators (Figure 2C). This organized functional specialization was disrupted in C1039G valves, where major ECM-synthetic and profibrotic genes became concentrated within the mVIC1 population (Figure 3B). In addition, mVIC1 cells from C1039G valves exhibited significantly increased expression of collagen-, elastin-, and GAG-biosynthetic genes, together with profibrotic genes (Figure 3C). Expression of *Tgfb1* and its receptors, *Tgfbr1* and *Tgfbr2*, was also significantly increased, supporting enhanced TGF-β pathway activation in mVIC1 from C1039G valves (Figure 3C and Supplemental Figure 3A).

Moreover, mVIC1 from C1039G valves displayed significantly increased senescence signature scores relative to WT control (Figure 3D and Supplemental Table 1), indicating activation of a senescence-associated transcriptional program. Compared with other VIC clusters, mVIC1 also exhibited marked enrichment of representative tricarboxylic acid (TCA) cycle and oxidative phosphorylation genes, consistent with increased mitochondrial metabolic engagement and a shift toward oxidative metabolic programming (Supplemental Figure 3B). This metabolic profile aligns with the heightened biosynthetic and remodeling demands of activated stromal cells (27,28). Notably, canonical myofibroblast markers were not enriched in mVIC1 but were preferentially expressed in other VIC clusters (Supplemental Figure 3C).

Integrating its activated ECM transcriptional profile, senescence-associated features, metabolic reprogramming, expansion in C1039G valves, and spatial enrichment at leaflet tips-the primary region affected in MMVD-we designated mVIC1 as activated mVIC (act-mVIC). Collectively, these features identify act-mVIC as a major disease-associated remodeling population associated with the pathological ECM changes characteristic of MMVD.

We next performed pseudotime trajectory analysis. VICs were ordered along a continuum progressing toward the act-mVIC state (Supplemental Figure 4A-B). Along this trajectory, expression of major ECM genes progressively increased, consistent with a transition toward a profibrotic phenotype. Although pseudotime analysis does not establish lineage hierarchy, these data are compatible with act-mVIC representing a terminally activated disease-associated state.

To assess regional gene expression patterns, spatial transcriptomic profiling was performed on mitral valve sections from C1039G and WT valves (21). Unsupervised clustering identified five major clusters corresponding to ventricular cardiomyocytes, atrial cardiomyocytes, blood cells, and two VIC populations (sp-mVIC1 and sp-mVIC2) (Figure 3E and Supplemental Figure 4C). Among these, sp-mVIC1 was markedly increased in C1039G valves and localized predominantly to the leaflet tip. Regions enriched for sp-mVIC1 in C1039G valves exhibited elevated expression of major ECM-related genes, including *Col1a1, Col3a1, Eln,* and *Has2* (Figure 3F). This transcriptional profile of sp-mVIC1 resembled that of act-mVIC identified by scRNA-seq, although the resolution of spatial transcriptomics precluded direct one-to-one mapping between spatial and scRNA-seq clusters.

Ligand-receptor interaction analysis revealed increased intercellular communication in C1039G valves compared with WT controls, predominantly driven by ECM-integrin signaling (Figure 3G) (19,29). A collagen IV-centered communication module was upregulated, with VIC-derived *Col4a1* exhibiting increased predicted interactions with matrix-sensing and mechanotransduction receptors, including *Itga1-Itgb1, Itgav-Itgb8, Cd44,* and *Sdc4*. These predicted interactions are consistent with enhanced integrin-dependent focal adhesion signaling and may facilitate force-mediated activation of latent TGF-β within a mechanically remodeled matrix microenvironment (30–32). *Col4a1*-associated signaling also extended to macrophages, where *Sdc4* and *Cd44* displayed elevated interaction strength, suggesting enhanced matrix sensing within collagen-enriched regions. In parallel, *Gas6-Axl* signaling was increased in VIC populations, consistent with activation of pro-survival and profibrotic transcriptional programs (33,34).

Together, these findings support a coordinated signaling framework in C1039G valves in which VIC-driven ECM remodeling, integrin-mediated mechanotransduction, and *Gas6-Axl* signaling converge to sustain the act-mVIC phenotype while promoting a matrix-dependent immunoregulatory niche that maintains macrophage engagement and reinforces a self-perpetuating profibrotic remodeling circuit characteristic of MMVD.

### Shared fibrotic remodeling and ECM disruption in human sporadic and Marfan- associated MVP

To determine whether findings from the murine C1039G model extend to human disease, we analyzed mitral valve specimens from three groups: normal controls, sporadic MVP, and Marfan-associated MVP.

Gross examination of normal mitral valves revealed thin, semitranslucent leaflets with smooth surfaces and well-defined free margins. The anterior and posterior leaflets were symmetric and pliable, with intact, slender chordae tendineae and preserved overall architecture. In contrast, sporadic and Marfan-associated MVP specimens exhibited leaflet thickening and enlargement, reduced translucency, and increased tissue firmness, consistent with fibrotic remodeling (Figure 4A). These gross features paralleled the structural abnormalities observed in C1039G valves. Because of current surgical practices, samples from sporadic MVP primarily included leaflet tips and certain mid-leaflet segments, whereas other valve specimens included entire leaflets. Care was taken to ensure that anatomically comparable regions were analyzed across normal and diseased samples.

**Figure 4.**
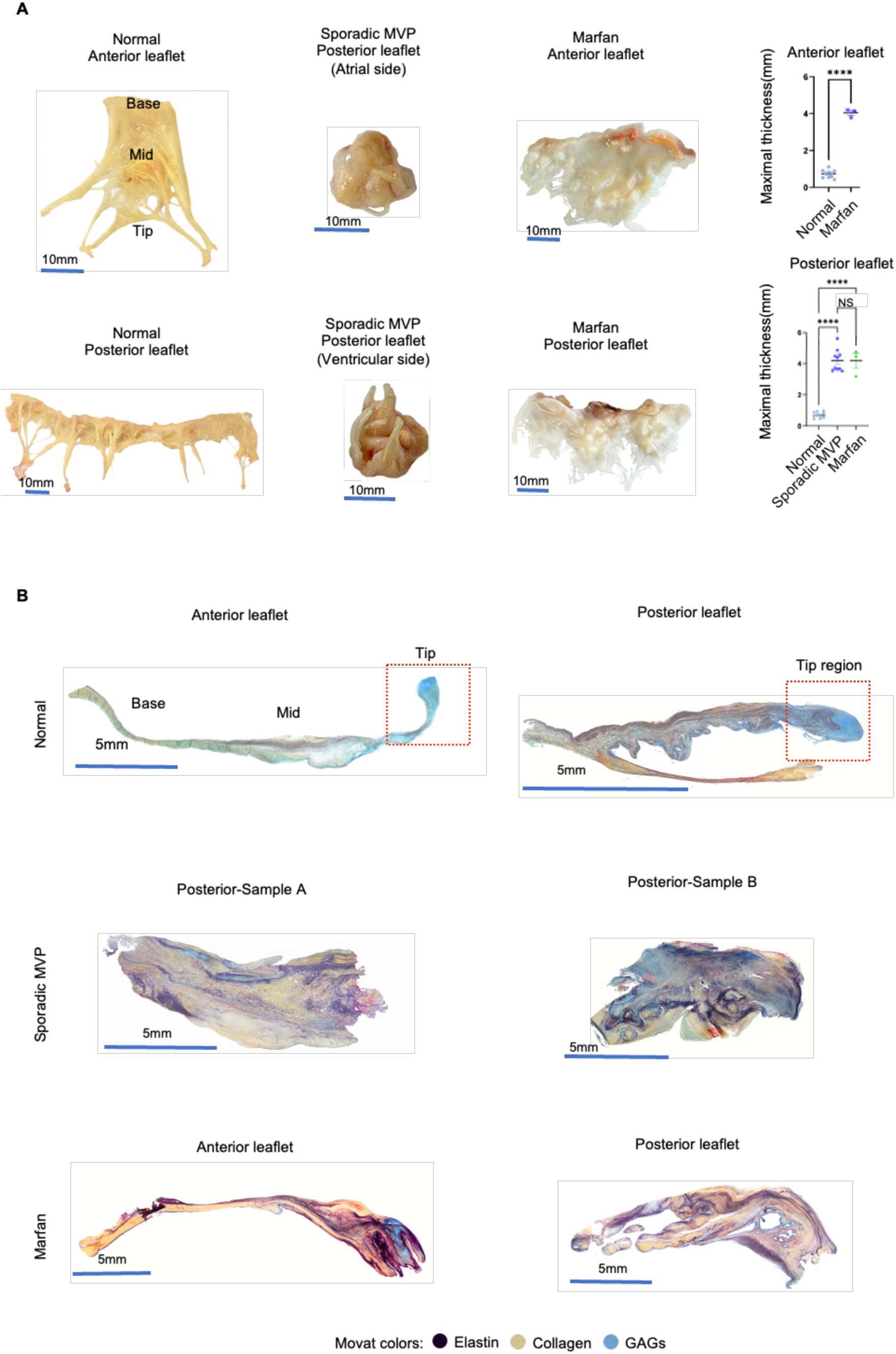
Structural and fibrotic features of sporadic and Marfan-associated mitral valve prolapse. (**A**) Gross morphology of mitral valve leaflets with quantification of maximal leaflet thickness. Sporadic MVP specimens consisted primarily of the leaflet tip, whereas other valve types included the full-length leaflet. Data are presented as individual data points with mean ± SEM. NS, not significant; ****P < 0.0001, by one-way ANOVA or unpaired t test. (**B**) Representative Movat’s pentachrome-stained histological sections illustrating ECM organization. The upper surface corresponds to the left atrial side, and the lower surface corresponds to the left ventricular side. Leaflet regions are labeled as tip, mid, and base.

Movat’s pentachrome staining was performed to assess ECM composition and architectural organization of collagen, elastin, and GAGs. The mitral leaflet is classically described as having a trilaminar structure (35). However, whole-leaflet Movat’s pentachrome staining revealed that the leaflet tip exhibited a more complex structural organization, characterized by an elastin-rich core encased by GAGs, forming a distinct “GAG-elastin sandwich.” The mid-leaflet and basal regions displayed a more conventional layered arrangement. In both sporadic and Marfan-associated MVP valves, this organized architecture was disrupted, with loss of normal layering, collagen-rich expansion, ECM disorganization, and leaflet thickening (Figure 4B).

Collectively, these gross and histologic analyses demonstrate that sporadic and Marfan-associated MVP share a common structural phenotype characterized by leaflet thickening, collagen-dominant fibrotic remodeling, and disruption of normal ECM organization, closely recapitulating the remodeling features observed in C1039G valves.

### Activated human VIC subset drives fibrotic remodeling in sporadic MVP

To further dissect the cellular heterogeneity underlying the observed structural and matricellular changes, we performed scRNA-seq on sporadic MVP valve specimens (Supplemental Table 2). Isolating viable single cells from human mitral leaflets was technically challenging because of the dense ECM architecture. After stringent quality-control filtering, 16,267 high-quality cells were retained for downstream analysis.

Unsupervised clustering followed by annotation using curated marker genes identified five major cell types (Figure 5A-B) (17,18), including VICs, endothelial cells, T cells, macrophages, and mast cells.

**Figure 5.**
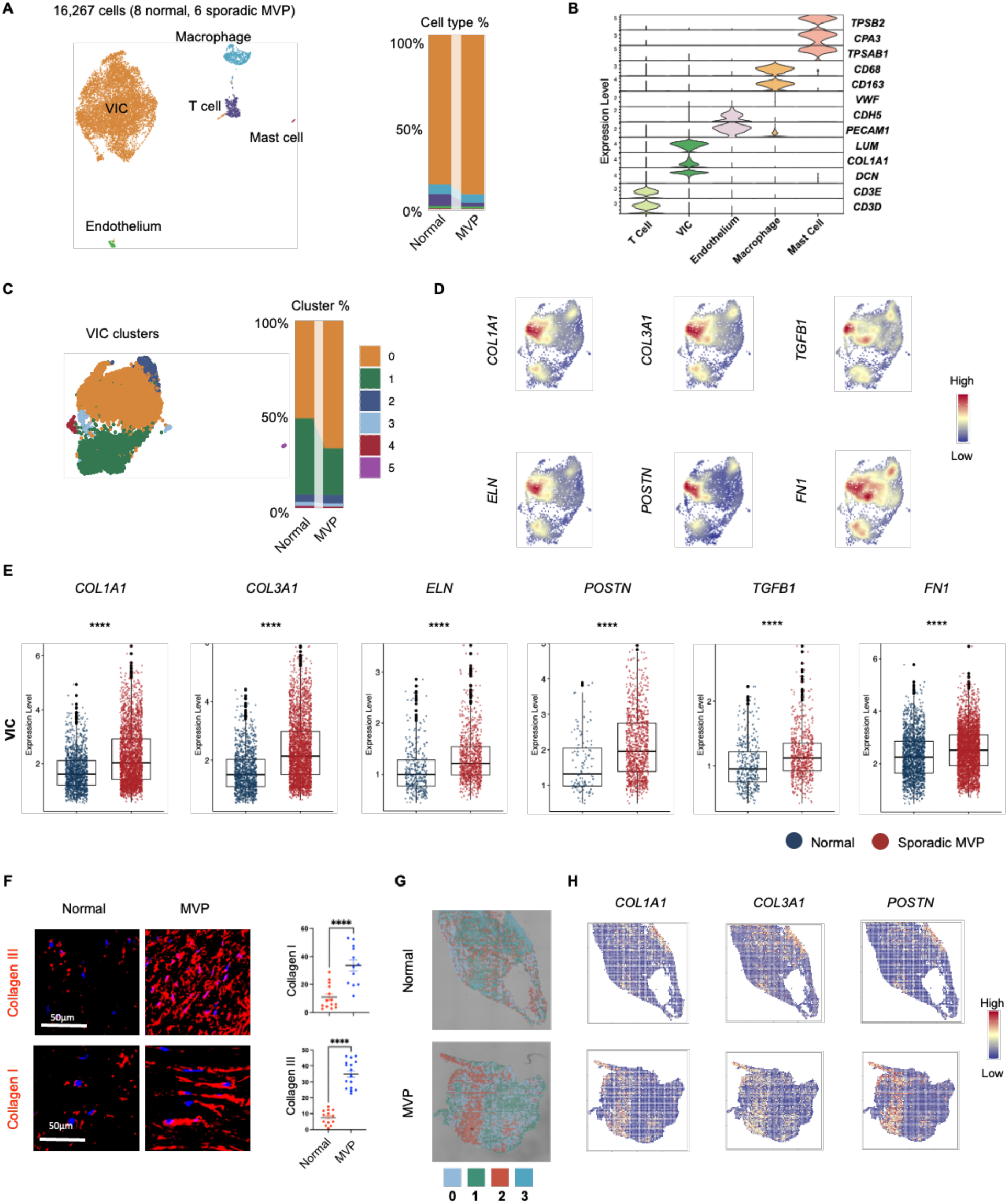
Integrated single-cell and spatial transcriptomic profiling of sporadic MVP. (**A**) UMAP visualization of normal and sporadic MVP mitral valves, with bar plots indicating the relative proportions of major cell types. (**B**) Violin plots showing expression of canonical marker genes across annotated valve cell populations. (**C**) UMAP showing unsupervised clustering of VICs. Bar plots indicate the relative proportions of each VIC cluster in normal and sporadic MVP valves. (**D**) UMAP demonstrating enrichment of representative profibrotic genes within a shared VIC cluster. (**E**) Boxplots quantifying expression of selected genes in the activated VIC cluster comparing normal and sporadic MVP valves. Data are shown as individual data points with mean ± SEM. Statistical significance was assessed using Wilcoxon test. ****P < 0.0001. (**F**) Representative immunofluorescence images of mitral valve sections stained for collagen I and collagen III, with corresponding quantification. Data are presented as individual data points with mean ± SEM. Statistical comparisons were performed using unpaired t test. ****P < 0.0001. (**G**) Spatial transcriptomic analysis showing unsupervised clustering of mitral valve sections. (**H**) Spatial expression patterns of representative genes related to fibrosis.

Unsupervised clustering of VICs revealed substantial transcriptional heterogeneity (Figure 5C and Supplemental Figure 5A). A distinct VIC subset (cluster 0) was characterized by marked enrichment of profibrotic and ECM-related genes, including *COL1A1, COL3A1, POSTN, ELN, FN1,* and *TGFB1* (Figure 5D). Cluster 0 was expanded in sporadic MVP relative to normal controls (Figure 5C) and displayed significantly higher expression of representative profibrotic genes compared with normal valves (Figure 5E).

Immunofluorescence staining for major collagens showed increased collagen deposition in sporadic MVP leaflets (Figure 5F) and RT-PCR analysis confirmed increased expression of ECM components and ECM modulators in MVP (Supplemental Figure 5B). Together, these findings identify a transcriptionally activated ECM-producing VIC population associated with human sporadic MVP.

To further characterize functional programs within VIC clusters, we performed gene set enrichment analysis (GSEA) using differentially expressed genes (DEGs) from sporadic MVP versus normal valves. ECM-related pathways were significantly enriched in sporadic MVP VICs (Supplemental Figure 6A). Metabolic pathway analysis revealed enrichment of oxidative phosphorylation and TCA cycle in sporadic MVP VICs, whereas glycolysis/gluconeogenesis-related signatures were relatively enriched in normal VICs (Supplemental Figure 6B). Although functional metabolic flux was not directly measured, this transcriptional pattern is consistent with increased mitochondrial oxidative metabolism in MVP VICs, potentially supporting sustained ECM biosynthesis and matrix remodeling demands.

Based on its coordinated ECM activation signature, metabolic reprogramming profile, and proportional expansion in sporadic MVP, this VIC population (cluster 0) was designated activated human VIC (act-hVIC).

To define regional transcriptional organization, spatial transcriptomic sequencing was performed on human mitral valve sections (Supplemental Table 3). Unsupervised clustering identified four major spatial clusters (Figure 5G). Profibrotic ECM-related genes were preferentially enriched in spatial cluster 2, which was expanded in sporadic MVP leaflets relative to normal controls (Figure 5H). This transcriptional profile resembled that of act-hVIC identified by scRNA-seq, although the resolution of spatial transcriptomics precluded direct one-to-one mapping between spatial and single-cell clusters.

### Profibrotic and inflammatory remodeling characteristics of sporadic MVP

Immunofluorescence staining demonstrated significant increased CD45⁺ immune cell infiltration in sporadic MVP valves compared with normal controls (Figure 6A), indicating enhanced inflammatory cell recruitment. Within macrophages, scRNA-seq analysis revealed significantly elevated expression of *TGFB1, TGFBR1*, and *TGFBR2* in sporadic MVP specimens (Figure 6B), indicating activation of macrophage-associated TGF-β signaling.

**Figure 6.**
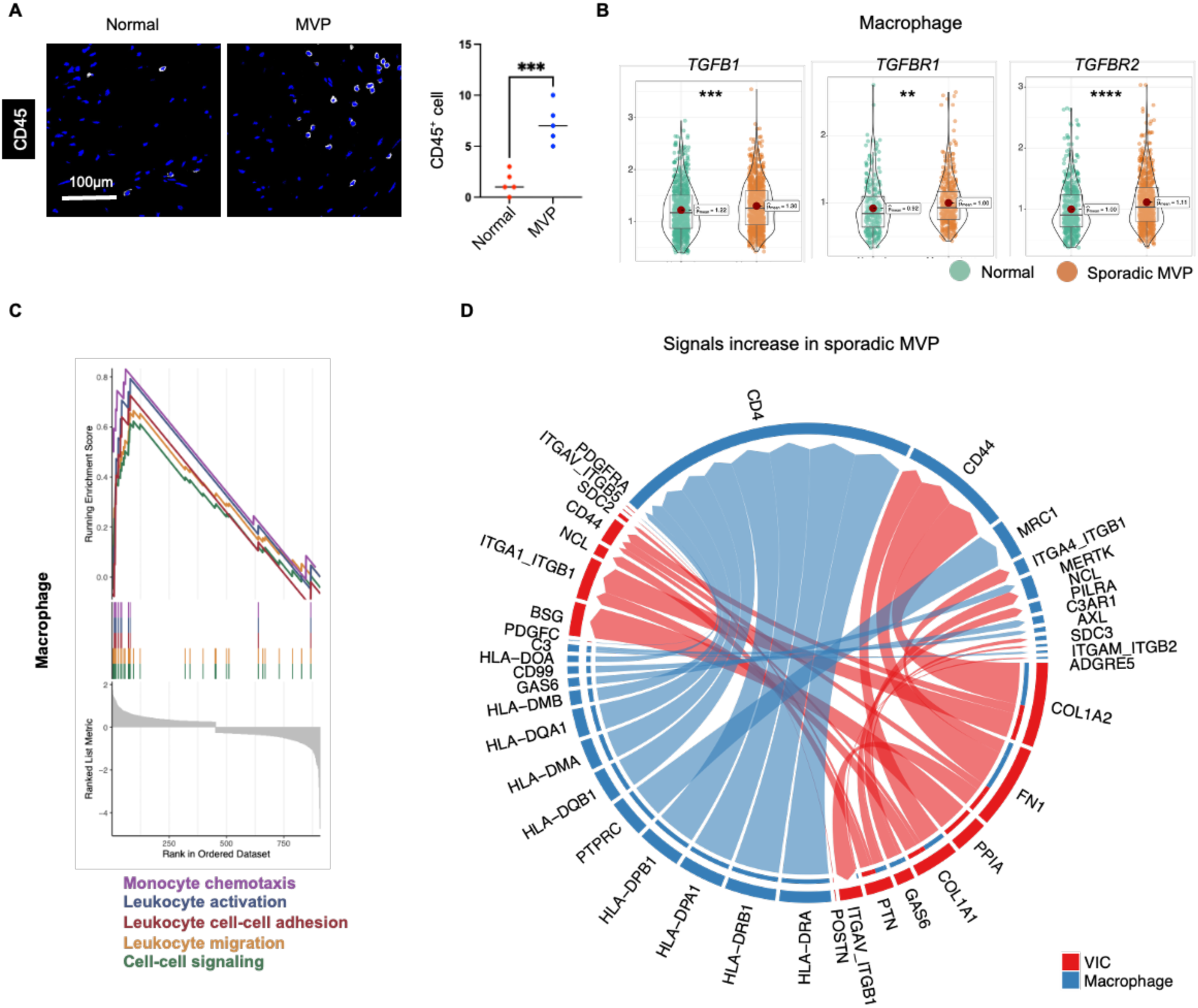
Inflammatory and profibrotic remodeling in sporadic MVP. **(A**) Representative immunofluorescence images of CD45 staining in normal and sporadic MVP mitral valve leaflets, with corresponding quantification. Data are presented as individual data points with mean ± SEM. Statistical comparisons were performed using unpaired t test. ***P < 0.001. (**B**) Violin plots comparing expression of representative profibrotic genes in macrophages from normal and sporadic MVP valves. Data are shown as individual data points with mean ± SEM. Statistical significance was assessed using Wilcoxon test. **P < 0.01; ***P < 0.001; ****P < 0.0001. (**C**) GSEA of DEGs in macrophages from sporadic MVP compared with normal mitral valves. (**D**) Representative ligand-receptor interaction networks upregulated in sporadic MVP compared with normal mitral valves. Edge thickness represents inferred interaction strength.

GSEA further demonstrated enrichment of pathways related to monocyte chemotaxis, leukocyte activation, cell-cell adhesion, and migration in sporadic MVP macrophages (Figure 6C), supporting an activated inflammatory phenotype.

Ligand-receptor interaction analysis demonstrated increased predicted communication between macrophages and VICs in sporadic MVP compared with normal valves (Figure 6D). ECM-integrin signaling was prominently enriched, including collagen-integrin pairs (*COL1A1/COL1A2-ITGA/ITGB*), as well as *FN1*- and *POSTN*-associated interactions. These findings are consistent with enhanced matrix-dependent adhesion and integrin-mediated mechanotransduction within profibrotic VIC populations. Notably, ECM ligand-integrin interactions also involved macrophages, suggesting increased matrix sensing and adhesion within collagen- and periostin-rich microenvironments.

Collectively, these findings indicate strengthened reciprocal ECM-centered communication between macrophages and VICs in sporadic MVP, consistent with establishment of a self-reinforcing profibrotic remodeling niche.

### Conserved profibrotic VIC subset is enriched in Marfan-associated MVP

We next performed scRNA-seq on human Marfan-associated MVP specimens (Supplemental Table 2). After stringent quality-control filtering, 12,901 high-quality cells were retained for downstream analysis. Annotation using curated marker genes identified the same five major cell populations observed in sporadic MVP (Figure 7A-B).

**Figure 7.**
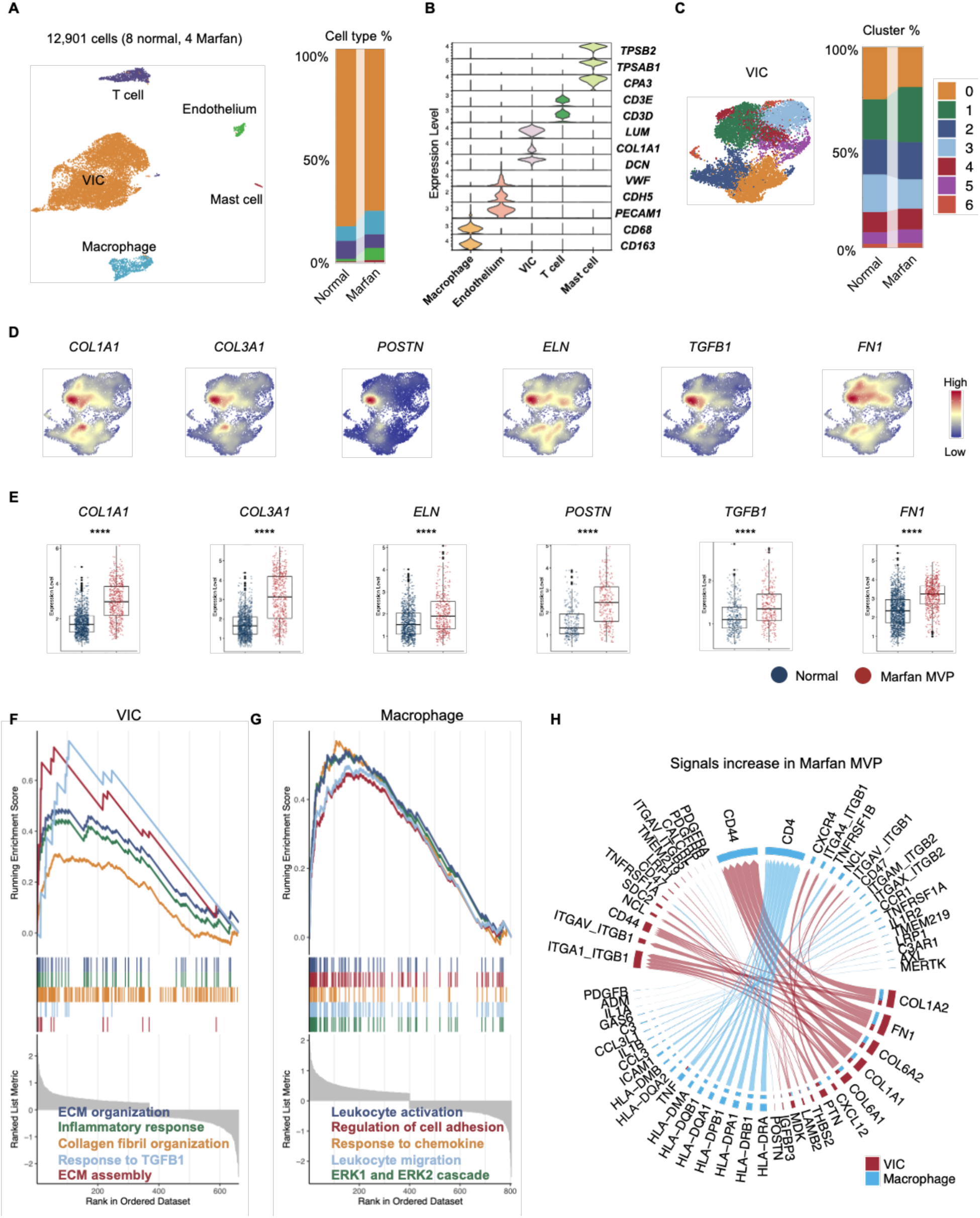
Single-cell transcriptomic analysis of Marfan-associated MVP. (**A**) UMAP visualization of scRNA-seq data from normal and Marfan-associated MVP mitral valves, with bar plots indicating the relative proportions of major cell types. (**B**) Violin plots showing expression of canonical marker genes across annotated valve cell populations. (**C**) UMAP showing unsupervised clustering of VICs, with bar plots indicating the relative proportions of each VIC cluster. (**D**) UMAP demonstrating enrichment of representative profibrotic genes within a shared activated VIC cluster. (**E**) Boxplots quantifying expression of selected genes in the activated human VIC cluster comparing normal and Marfan-associated MVP valves. Data are presented as individual data points with mean ± SEM. Statistical significance was assessed using Wilcoxon test. ****P < 0.0001. GSEA of DEGs in the activated VIC cluster (**F**) and macrophages (**G**) from Marfan-associated MVP compared with normal mitral valves. (**H**) Representative ligand-receptor interaction networks upregulated in Marfan-associated MVP relative to normal mitral valves. Edge thickness represents inferred interaction strength.

Unsupervised clustering of VICs revealed marked transcriptional heterogeneity (Figure 7C and Supplemental Figure 7A). A distinct VIC subset (cluster 1) demonstrated preferential enrichment of profibrotic and ECM-associated genes and was markedly expanded in Marfan valves compared with normal controls (Figure 7C-D). Moreover, this cluster exhibited significantly higher expression of these profibrotic genes in Marfan valves relative to normal valves (Figure 7E).

GSEA of DEGs from Marfan-associated MVP versus normal valves within this VIC cluster demonstrated significant enrichment of pathways related to ECM organization and extracellular structure remodeling (Figure 7F), consistent with activation of a conserved matrix-remodeling program. Metabolic pathway analysis further revealed enrichment of oxidative phosphorylation and TCA cycle pathways in Marfan-associated MVP, whereas glycolysis and gluconeogenesis signatures were relatively enriched in normal valves (Supplemental Figure 7B).

Based on its coordinated ECM activation signature, metabolic reprogramming profile, and proportional expansion in Marfan-associated MVP, this VIC population (cluster 1) was designated act-hVIC, consistent with the activated VIC population identified in sporadic MVP.

Macrophages from Marfan-associated MVP valves exhibited significant enrichment of pathways related to leukocyte activation, chemokine response, regulation of cell adhesion, leukocyte migration, and ERK1/ERK2 signaling compared with normal controls (Figure 7G). These transcriptional programs are consistent with an activated inflammatory phenotype characterized by enhanced migratory capacity and engagement of MAPK-associated signaling pathways (36).

Ligand-receptor interaction analysis demonstrated markedly increased predicted signaling interactions between macrophages and VICs in Marfan-associated MVP compared with normal valves (Figure 7H). Enhanced communication was predominantly driven by ECM-integrin axes, including collagen-integrin (*COL1A1/COL1A2-ITGA/ITGB*) and *FN1*-integrin interactions. These predicted interactions are compatible with intensified matrix-dependent adhesion and integrin-mediated mechanotransduction within remodeled leaflet regions (29).

Collectively, these findings indicate amplified reciprocal communication between profibrotic VICs and activated macrophages in Marfan valves, consistent with reinforcement of a fibro-inflammatory remodeling microenvironment.

### Conserved activated VIC program is shared across human and murine MMVD

To directly compare transcriptional programs across species and disease etiologies, we integrated scRNA-seq datasets from human mitral valves (normal, sporadic MVP, and Marfan-associated MVP) and murine MMVD samples. We first integrated datasets from human normal, sporadic MVP, and Marfan-associated MVP valves (Figure 8A). Using curated marker genes, the same five major cell populations were consistently identified (Figure 8B-C).

**Figure 8.**
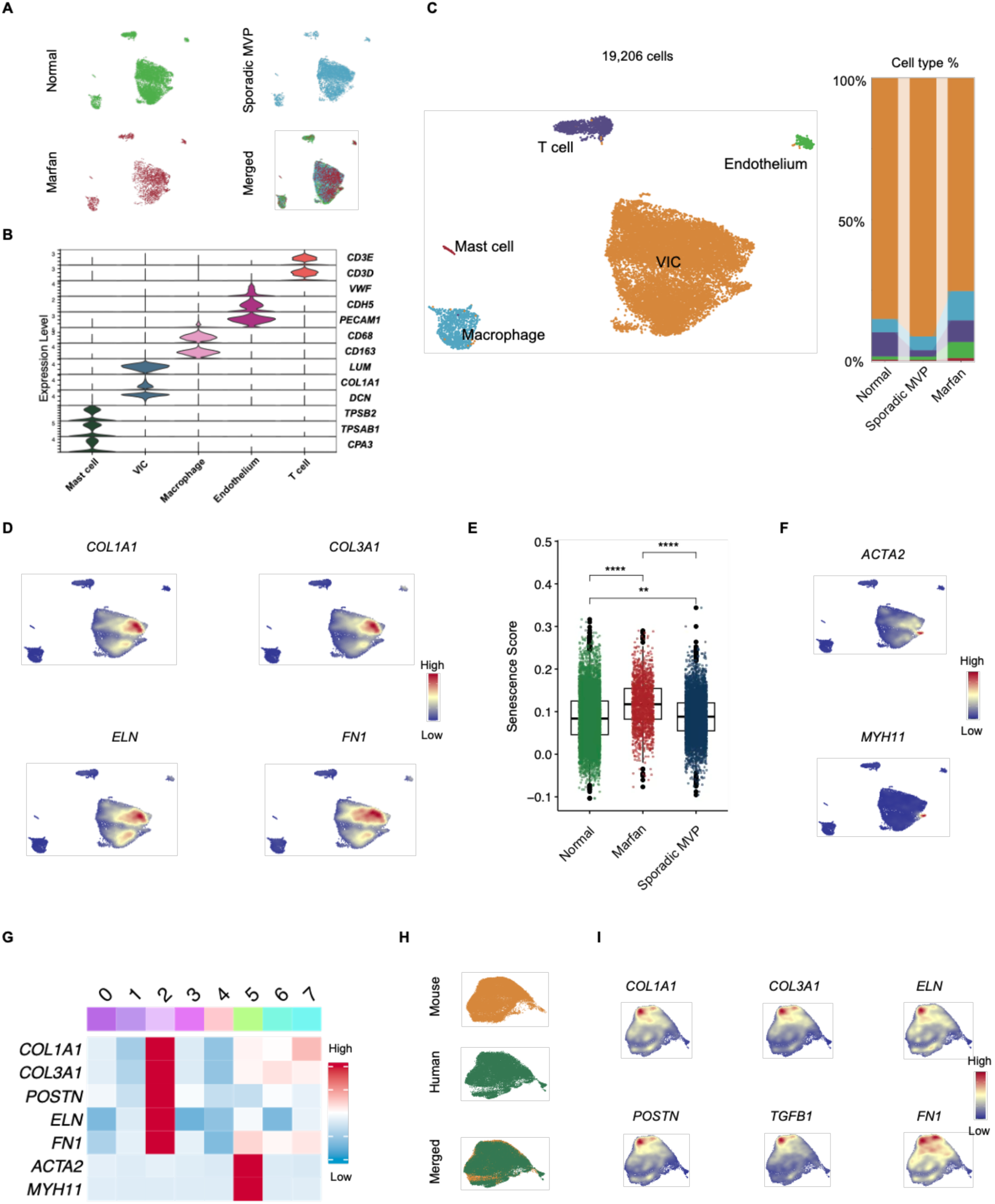
Cross-species integration showing conserved profibrotic VIC programs in mitral valve disease. (**A**) UMAP visualization of integrated scRNA-seq data from normal, sporadic MVP, and Marfan-associated MVP mitral valves. (**B**) Violin plots depicting expression of canonical marker genes across annotated valve cell populations. (**C**) UMAP highlighting major cell types, with bar plots indicating the relative proportions of each cell population. (**D**) UMAP demonstrating enrichment of representative profibrotic genes within a shared VIC cluster. (**E**) Boxplot comparing SenMayo senescence signature scores in the activated human VICs. Data are presented as individual data points with mean ± SEM. Statistical significance was assessed using Wilcoxon test. **P < 0.01; ****P < 0.0001. (**F**) UMAP showing expression of canonical myofibroblast markers across VIC populations. (**G**) Heatmap displaying representative gene expression patterns across VIC clusters. (**H**) Cross-species UMAP integration of VICs from mouse and human mitral valves. (**I**) UMAP demonstrating enrichment of representative profibrotic genes within a shared VIC cluster cross-species.

Fibrotic ECM-related genes, including *COL1A1, COL3A1, ELN*, and *FN1* were enriched within a distinct VIC population (Figure 8D), corresponding to the previously defined activated, profibrotic VIC subset. Notably, act-hVIC populations identified in both sporadic and Marfan-associated MVP localized to the same cluster after integration, indicating convergence on a shared transcriptional state across disease etiologies.

This fibrotic VIC population exhibited significantly increased senescence signature scores in diseased valves compared with normal controls (Figure 8E and Supplemental Table 1), indicating enrichment of a senescence-associated transcriptional program within the activated VIC compartment.

Notably, canonical myofibroblasts markers such as *ACTA2* and *MYH11* were enriched within a separate VIC population and did not broadly overlap with this fibrotic population (Figure 8F). Additional unsupervised clustering of VICs (Supplemental Figure 8) further demonstrated segregation between a profibrotic ECM-enriched cluster 2 and a distinct cluster 5 enriched for canonical myofibroblast markers (Figure 8G).

Collectively, these findings indicate that disease-associated act-hVICs represent a transcriptionally activated, profibrotic ECM-producing state that is molecularly distinct from classical myofibroblast populations.

Given leaflet thickening and increased mechanical stiffness observed in both sporadic and Marfan-associated MVP valves, we evaluated whether pathological calcification contributes to disease pathogenesis. Von Kossa and Alizarin Red staining failed to demonstrate mineral deposition in sporadic or Marfan-associated mitral valves (Supplemental Figure 9). High-resolution micro-computed tomography (micro-CT) also did not detect calcified regions within diseased leaflets (Supplemental Video 1-4). These findings indicate that leaflet stiffening in MMVD occurs in the absence of overt calcification and is instead attributable primarily to fibrotic ECM remodeling and matrix reorganization. It should be noted, however, that patients with evident calcification are typically excluded from mitral valve repair surgery and therefore not present in our surgical specimen. Although this does not exclude a role for calcification in mitral valve diseases more broadly, our findings suggest that ECM components-particularly collagen deposition and fibrosis-play a significant role in mitral valve disease progression.

To assess cross-species conservation, we next integrated human and murine VIC datasets (Figure 8H). Representative profibrotic genes were consistently enriched within a corresponding VIC population across species (Figure 8I), demonstrating strong cross-species conservation of this activated transcriptional program. The human act-hVIC population aligned transcriptionally with the murine act-mVIC cluster identified in the Fbn1-deficient model, supporting the presence of a shared disease-associated VIC state across sporadic MVP, Marfan-associated MVP, and murine MMVD.

Together, these findings establish that MMVD is characterized by a conserved activated VIC program that transcends species and genetic etiology. Discussion

In this study, we constructed a cross-species, multimodal atlas of normal and diseased mitral valves by integrating single-cell and spatial transcriptomics with comprehensive histopathology in both murine model and human sporadic and Marfan-associated MVP. Across species, we identified a conserved activated VIC subset enriched within mechanically vulnerable leaflet tip regions. Rather than representing diffuse global VIC activation, this subset forms a spatially restricted ECM-centered signaling module characterized by enhanced matrix production and intensive ECM-integrin-mediated crosstalk with macrophages. These findings refine the conceptual framework of mitral valve disease as a spatially organized, VIC-driven network disorder.

Because of slender and elongated structure of the mitral leaflet, its full structural complexity has historically been simplified into a trilaminar description (35). Whole-leaflet Movat’s pentachrome staining instead reveals a more regionally specialized organization aligned with biomechanical demands. The leaflet base contains two functional layers: an upper elastin-rich layer (atrial side) provides elasticity while a lower collagen-rich layer (ventricular side) confers tensile strength. The mid-leaflet transitions into a tri-layered configuration with an elastin core flanked by collagen layers, balancing flexibility and mechanical resilience. The leaflet tip contains a GAG-enriched domain enveloping an elastin core, forming a compressible yet resilient structure that maintains load-bearing capacity while preserving elastic recoil under repetitive mechanical stress. In diseased valves, VIC activation preferentially occurs within this normally GAG-rich region, leading to excessive ECM production, collagen accumulation, architectural distortion, and immune cell recruitment. These alterations disrupt matrix stratification and mechanical homeostasis, thereby promoting progressive leaflet thickening.

Although MMVD has historically been described as a “myxomatous” disease, the contribution of fibrosis-associated ECM remodeling to disease progression has remained insufficiently defined. This knowledge gap partly reflects the technical challenges of systematically profiling the mitral leaflet, whose thin and elongated structure complicates comprehensive spatial analysis. A central finding of the present study is the identification of fibrotic ECM remodeling as a prominent pathological feature of MMVD. These data suggest that fibrosis represents an important structural and mechanistic component of mitral valve disease beyond the classical myxomatous description.

A significant finding of this study is the identification of an activated VIC state that diverges from the canonical TGF-β-driven myofibroblast paradigm. Classical models emphasize induction of contractile genes such as *ACTA2, MYH11*, and *CNN1*. In contrast, the act-VIC population in both species shows strong enrichment for profibrotic ECM synthesis genes (*COL1A1, COL3A1, POSTN,* and *FN1*) and matrix remodeling programs without consistent upregulation of canonical myofibroblast markers (37). This transcriptional configuration resembles matrifibrocyte-like states described in chronic cardiac fibrosis, characterized by sustained ECM production rather than contractility activation (37). Although definitive lineage relationships will require formal validation, these data suggest that chronic mitral valve remodeling is mediated primarily by a persistent ECM-producing VIC state rather than classical myofibroblast differentiation.

Therapeutically, this distinction suggests that targeting ECM production, matrix-receptor interactions, or inflammatory signaling may be more effective than strategies aimed solely at suppressing classical myofibroblast differentiation.

Our findings further indicate that act-VICs exhibit coordinated senescence-associated and metabolic transcriptional features (38–40). SenMayo signature scores were selectively elevated in act-VIC populations across murine and human datasets, supporting engagement of a senescence-like program. In parallel, pathway enrichment analysis demonstrated increased representation of TCA cycle and oxidative phosphorylation signatures. Although metabolic flux was not directly assessed, this transcriptional enrichment is consistent with enhanced mitochondrial metabolic engagement, potentially supporting the energetic and biosynthetic demands of sustained ECM synthesis and matrix remodeling. Emerging evidence from fibrotic disorders indicates that mitochondrial metabolic reprogramming can reinforce profibrotic transcriptional states (41,42). Together, these findings suggest that VIC activation involves coordinated transcriptional, metabolic, and senescence-associated remodeling programs that may stabilize a persistent pathogenic state in MMVD. These observations further raise the possibility that targeting metabolic or senescence-associated pathways may have therapeutic value.

Ligand-receptor inference indicates that VIC-macrophage communication is prominently enriched for ECM-integrin interactions. Collagen-integrin pairs (*COL1A1/COL1A2-ITGA1/ITGB1, COL4A1-ITGA1/ITGB1*), periostin-integrin (*POSTN-ITGA5/ITGB5*), and thrombospondin-CD47 (*THBS-CD47*) axes were prominently enriched, along with additional integrin-associated signaling pathways (43–45). These interactions provide a mechanistic framework linking matrix remodeling to mechanotransduction and immune activation. As act-VICs deposit collagen and other ECM components, they reshape the mechanical and structural properties and ligand landscape of the leaflet microenvironment, thereby enhancing integrin signaling in macrophages and other stromal cells. Activated macrophages, in turn, produce cytokines, chemokines, and profibrotic mediators that further amplify VIC activation and ECM deposition (46,47). This establishes a feed-forward loop in which ECM remodeling and immune activation reinforce one another, providing a mechanistic explanation for progressive leaflet thickening in both genetic and sporadic disease contexts.

Spatial mapping at transcriptomic and protein levels further demonstrated that pathogenic remodeling is regionally concentrated, particularly at leaflet tip regions rather than uniformly distributed across the whole leaflet. In murine valves, act-mVICs localize to thickened tip regions enriched for *Col1a1, Col3a1*, *Eln*, and *Has2*. In human valves, act-hVIC clusters expanded within fibrotic regions interspersed with macrophage infiltrates. These findings suggest that intrinsic regional heterogeneity, together with localized mechanical stress (48,49), predisposes specific leaflet domains to ECM degeneration. Disease progression therefore appears to arise from amplification of regionally restricted signaling modules rather than uniform activation across the leaflet.

Across both species, a shared core signature emerges characterized by expansion of a distinct profibrotic VIC population, increased macrophage abundance with inflammatory and ECM-remodeling transcriptional profiles, and intensified VIC-macrophage communication through ECM-integrin signaling axes. These conserved features indicate that the act-VIC population represents a cross-species pathogenic module operative in both genetic and non-syndromic mitral valve disease. From a translational perspective, these data support the C1039G model as a mechanistically relevant platform for interrogating VIC- and macrophage-targeted interventions with potential applicability to human MMVD.

Several limitations should be considered. First, upstream etiologies and remodeling kinetics differ between species. The C1039G model represents a genetically driven and relatively rapid remodeling process, whereas human MVP progresses over years to decades. Chronic human disease likely involves additional ECM maturation, immune adaptation, and cumulative mechanobiological remodeling that may not be fully recapitulated in the murine system. Moreover, matrix composition differs between species, with murine valves demonstrating coordinated increases in collagen, elastin, and GAGs, whereas human MVP specimens show more collagen-dominant deposition. These differences should be acknowledged despite the shared transcriptional program. Second, human tissue availability imposes important constraints. Surgical mitral valve specimens are limited to patients undergoing operative intervention and therefore may not fully represent the broader clinical spectrum of MVP, particularly early or subclinical stages. The current cohort also capture substantial biological heterogeneity across patients, including differences in demographics, disease duration, and remodeling stage. Because human valve remodeling is likely asynchronous, with multiple disease phases coexisting within individual specimens, human VIC subsets may not segregate into sharply defined functional states as observed in the murine model. Instead, their transcriptional programs may exist along a continuum, making subtype-specific functional assignments less distinct. Larger, stage-stratified studies with expanded demographic representation and longitudinal sampling will be necessary to determine how act-VIC dynamics, immune infiltration, and ECM-integrin signaling networks evolve over time. Third, spatial transcriptomic coverage in human valves was necessarily restricted to pathologically relevant regions because of platform size limitations, and region-specific programs outside the leaflet tip may therefore be underrepresented.

Fourth, spatial transcriptomic resolution does not achieve true single-cell assignment. Fifth, trajectory inference approaches cannot establish definitive lineage relationships; in vivo lineage tracing will be required to determine whether act-VICs arise from specific baseline VIC subtypes or represent a distinct lineage.

In summary, integration of single-cell and spatial transcriptomic datasets across murine and human mitral valves identifies a conserved activated VIC state that localizes to mechanically vulnerable leaflet tip regions and orchestrates profibrotic remodeling programs. These act-VICs engage in intensive ECM-integrin-mediated crosstalk with macrophages, forming a self-reinforcing matrix-remodeling circuit that drives MMVD. These findings refine the current paradigm of mitral valve disease from diffuse, nonspecific myofibroblast activation toward a spatially organized, ECM-high VIC-centered pathogenic network. The data nominate fibrosis-associated transcriptional programs, chemokine signaling, integrin-dependent mechanotransduction, senescence-associated pathways, and mitochondrial metabolic remodeling as candidate therapeutic targets. Collectively, this cross-species analysis provides a mechanistic foundation and translational framework for the development of therapeutic interventions in MMVD.

## Author contributions

YL, RF, and AG designed the study. FG, MD, YL, DZ, XL, MY and MZ conducted experiments and acquired data. FG, MD, DZ, XL, MY, and YL analyzed RNA-seq data. FG, IM, IH wrote and edited the manuscript. GF, YL, RF, GT, MK, and AG analyzed and interpreted data, wrote and edited the manuscript.

## Acknowledgements

This work was supported by Yale’s Department of Surgery William W.L. Glenn endowed research fund. GF was partially supported by NIH R01 HL131872. YL was supported by NIH R35 GM150838 and NIH R01 HL173271. RF was supported by NIH UG3CA257393, UH3CA257393, U54AG076043, and U54AG079759. Abdulrahman Hassab at Yale School of Medicine assisted with collection of clinical tissue samples and clinical data.

## References

1. Delling FN, Vasan RS. Epidemiology and pathophysiology of mitral valve prolapse: new insights into disease progression, genetics, and molecular basis. Circulation. 2014;129:2158–70.

2. Freed LA, Levy D, Levine RA et al. Prevalence and clinical outcome of mitral-valve prolapse. N Engl J Med. 1999;341:1–7.

3. Narula N, Schwartz Y, Devereux RB et al. Clinical and phenotypic correlates of mitral valve prolapse in Marfan syndrome: The Cornell Aortic Aneurysm Registry. Journal of the American Heart Association. 2025;14:e040947.

4. Levine RA, Hagege AA, Judge DP et al. Mitral valve disease morphology and mechanisms. Nat Rev Cardiol .2015;12:689–710.

5. Roberts WC, Vowels TJ, Ko JM, Hebeler RF, Jr. Gross and histological features of excised portions of posterior mitral leaflet in patients having operative repair of mitral valve prolapse and comments on the concept of missing (= ruptured) chordae tendineae. J Am Coll Cardiol. 2014;63:1667–74.

6. Liu AC, Joag VR, Gotlieb AI. The emerging role of valve interstitial cell phenotypes in regulating heart valve pathobiology. Am J Pathol. 2007;171:1407–18.

7. Xu K, Xie S, Huang Y et al. Cell-type transcriptome atlas of human aortic valves reveal cell heterogeneity and endothelial to mesenchymal transition involved in calcific aortic valve disease. Arterioscler Thromb Vasc Biol. 2020;40:2910–2921.

8. Hulin A, Hortells L, Gomez-Stallons MV et al. Maturation of heart valve cell populations during postnatal remodeling. Development. 2019;146(12):dev173047.

9. Shu S, Fu M, Chen X et al. Cellular Landscapes of Nondiseased Human Cardiac Valves From End-Stage Heart Failure-Explanted Heart. Arterioscler Thromb Vasc Biol. 2022;42:1429–1446.

10. Mahler GJ, Butcher JT. Inflammatory regulation of valvular remodeling: the good(?), the bad, and the ugly. Int J Inflam. 2011;2011:721419.

11. Mahimkar R, Nguyen A, Mann M et al. Cardiac transgenic matrix metalloproteinase-2 expression induces myxomatous valve degeneration: a potential model of mitral valve prolapse disease. Cardiovasc Pathol. 2009;18:253–61.

12. Xu N, Yutzey KE. Therapeutic CCR2 blockade prevents inflammation and alleviates myxomatous valve disease in Marfan syndrome. JACC Basic Transl Sci. 2022;7:1143–1157.

13. Walker GA, Masters KS, Shah DN et al. Valvular myofibroblast activation by transforming growth factor-beta: implications for pathological extracellular matrix remodeling in heart valve disease. Circ Res. 2004;95:253–60.

14. Rabkin E, Aikawa M, Stone JR et al. Activated interstitial myofibroblasts express catabolic enzymes and mediate matrix remodeling in myxomatous heart valves. Circulation. 2001;104:2525–32.

15. Ng CM, Cheng A, Myers LA et al. TGF-beta-dependent pathogenesis of mitral valve prolapse in a mouse model of Marfan syndrome. J Clin Invest. 2004;114:1586–92.

16. Li G, Wang M, Caulk AW et al. Chronic mTOR activation induces a degradative smooth muscle cell phenotype. J Clin Invest. 2020;130:1233–1251.

17. Litviňuková M, Talavera-López C, Maatz H et al. Cells of the adult human heart. Nature. 2020;588:466–472.

18. Kuppe C, Ramirez Flores RO, Li Z et al. Spatial multi-omic map of human myocardial infarction. Nature. 2022;608:766–777.

19. Jin S, Guerrero-Juarez CF, Zhang L et al. Inference and analysis of cell-cell communication using CellChat. Nat Commun. 2021;12:1088.

20. Saul D, Kosinsky RL, Atkinson EJ et al. A new gene set identifies senescent cells and predicts senescence-associated pathways across tissues. Nat Commun. 2022;13:4827.

21. Liu Y, DiStasio M, Su G et al. High-plex protein and whole transcriptome co-mapping at cellular resolution with spatial CITE-seq. Nat Biotechnol. 2023;41:1405–1409.

22. Liu Y, Yang M, Deng Y et al. High-spatial-resolution multi-omics sequencing via deterministic barcoding in tissue. Cell. 2020;183:1665–1681.e18.

23. Navarro JF, Sjöstrand J, Salmén F et al. Pipeline: an automated pipeline for spatial mapping of unique transcripts. Bioinformatics 2017;33:2591–2593.

24. Stuart T, Butler A, Hoffman P et al. Comprehensive integration of single-cell data. Cell. 2019;177:1888–1902.e21.

25. Judge DP, Biery NJ, Keene DR et al. Evidence for critical contibution of haploinsufficiency in the complex pathogenesis of Marfan syndrome. J Clin Invest. 2004;114:172–181.

26. Kim AJ, Xu N, Umeyama K et al. Deficiency of circulating monocytes ameliorates the progression of myxomatous valve degeneration in Marfan syndrome. Circulation. 2020;141:132–146.

27. Para R, Romero F, George G et al. Metabolic reprogramming as a driver of fibroblast activation in pulmonary fibrosis. Am J Med Sci. 2019;357:394–398.

28. Li M, Riddle S, Zhang H et al. Metabolic reprogramming regulates the proliferative and inflammatory phenotype of adventitial fibroblasts in pulmonary hypertension through the transcriptional corepressor C-Terminal Binding Protein-1. Circulation. 2016;134:1105–1121.

29. Gao F, Chen Q, Mori M et al. Integrin-mediated mTOR signaling drives TGF-β overactivity and myxomatous mitral valve degeneration in hypomorphic fibrillin-1 mice. J Clin Invest. 2025;135(14):e183558.

30. Sarrazy V, Koehler A, Chow ML et al. Integrins αvβ5 and αvβ3 promote latent TGF-β1 activation by human cardiac fibroblast contraction. Cardiovasc Res. 2014;102:407–417.

31. Sheppard D. Integrin-mediated activation of latent transforming growth factor beta. Cancer Metastasis Rev. 2005;24:395–402.

32. Sun Z, Guo SS, Fässler R. Integrin-mediated mechanotransduction. J Cell Biol. 2016;215:445–456.

33. Espindola MS, Habiel DM, Narayanan R et al. Targeting of TAM receptors ameliorates fibrotic mechanisms in idiopathic pulmonary fibrosis. Am J Respir Crit Care Med 2018;197:1443–1456.

34. Bárcena C, Stefanovic M, Tutusaus A et al. Gas6/Axl pathway is activated in chronic liver disease and its targeting reduces fibrosis via hepatic stellate cell inactivation. J Hepatol. 2015;63:670–678.

35. Small AM, Yutzey KE, Binstadt BA et al. Unraveling the mechanisms of valvular heart disease to identify medical therapy targets: A Scientific Statement From the American Heart Association. Circulation. 2024;150:e109–e128.

36. Hulin A, Anstine LJ, Kim AJ et al. Macrophage transitions in heart valve development and myxomatous valve disease. Arterioscler Thromb Vasc Biol. 2018;38:636–644.

37. Amrute JM, Luo X, Penna V et al. Targeting immune–fibroblast cell communication in heart failure. Nature. 2024;635:423–433.

38. Tang Q, Tang K, Markby GR et al. Autophagy regulates cellular senescence by mediating the degradation of CDKN1A/p21 and CDKN2A/p16 through SQSTM1/p62-mediated selective autophagy in myxomatous mitral valve degeneration. Autophagy. 2025;21:1433–1455.

39. Tang Q, Markby GR, MacNair AJ et al. TGF-β-induced PI3K/AKT/mTOR pathway controls myofibroblast differentiation and secretory phenotype of valvular interstitial cells through the modulation of cellular senescence in a naturally occurring in vitro canine model of myxomatous mitral valve disease. Cell Prolif. 2023;56:e13435.

40. Grzeczka A, Graczyk S, Kordowitzki P. Involvement of TGF-β, mTOR, and inflammatory mediators in aging alterations during myxomatous mitral valve disease in a canine model. Geroscience. 2025;47:5401–5433.

41. Li Q, Larouche-Lebel É, Loughran KA et al. Metabolomics analysis reveals deranged energy metabolism and amino acid metabolic reprogramming in dogs with myxomatous mitral valve disease. J Am Heart Assoc. 2021;10:e018923.

42. Granata S, Bernardini C, Glogowski PA et al. Mitochondrial bioenergetic dysfunction linked to myxomatous mitral valve degeneration explored by PBMCs metabolism analysis. Biochim Biophys Acta Bioenerg. 2024;1865:149505.

43. Xu Y, Gurusiddappa S, Rich RL et al. Multiple binding sites in collagen type I for the integrins alpha1beta1 and alpha2beta1. J Biol Chem. 2000;275:38981–9.

44. Sanders MA, Basson MD. Collagen IV regulates Caco-2 migration and ERK activation via alpha1beta1- and alpha2beta1-integrin-dependent Src kinase activation. Am J Physiol Gastrointest Liver Physiol. 2004;286:G547–57.

45. Becker HM, Rullo J, Chen M et al. α1β1 integrin-mediated adhesion inhibits macrophage exit from a peripheral inflammatory lesion. J Immunol. 2013;190:4305–14.

46. Simões FC, Cahill TJ, Kenyon A et al. Macrophages directly contribute collagen to scar formation during zebrafish heart regeneration and mouse heart repair. Nat Commun. 2020;11:600.

47. Buechler MB, Fu W, Turley SJ. Fibroblast-macrophage reciprocal interactions in health, fibrosis, and cancer. Immunity. 2021;54:903–915.

48. Rausch MK, Bothe W, Kvitting JP et al. In vivo dynamic strains of the ovine anterior mitral valve leaflet. J Biomech. 2011;44:1149–57.

49. Krishnamurthy G, Itoh A, Bothe W et al. Stress-strain behavior of mitral valve leaflets in the beating ovine heart. J Biomech. 2009;42:1909–16.

