## Supplemental Files for "Cross-Species Multi-Omics Profiling Identifies Conserved Activated Valvular Interstitial Cell Population Driving Myxomatous Mitral Valve Degeneration"

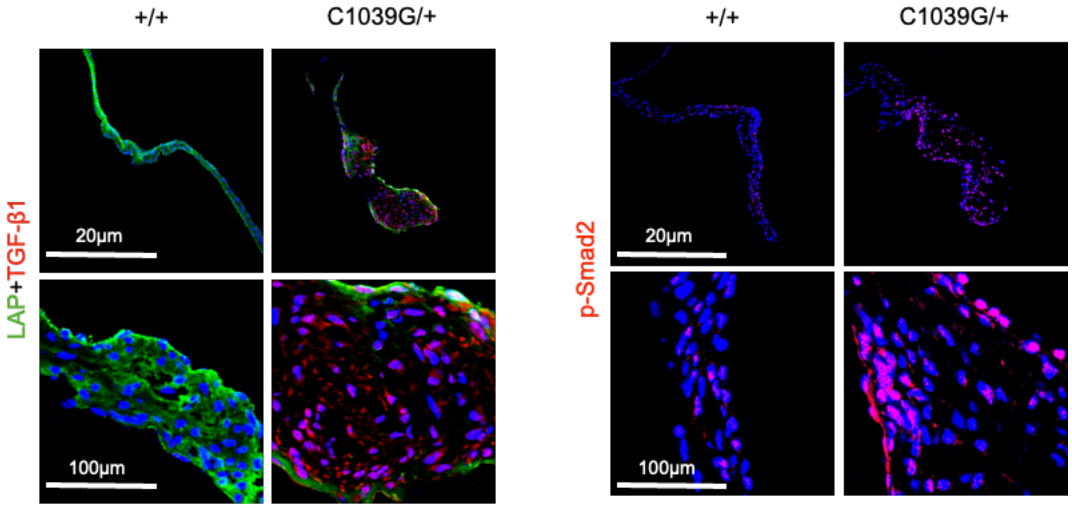


**Supplemental Figure 1. Characterization of mitral valve remodeling in Fbn1-deficient mice.** Representative immunofluorescence images of mitral valve sections stained for TGF-β1, latency-associated peptide (LAP), and phosphorylated Smad2 (p-Smad2).

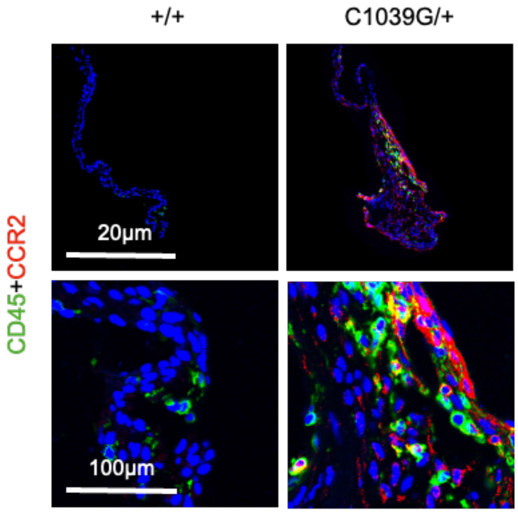


**Supplemental Figure 2. Characterization of inflammatory remodeling in Fbn1-deficient mice.** Representative immunofluorescence images of mitral valve sections stained for CD45 and CCR2.

**Supplemental Figure 3 (related to Figure 3)**

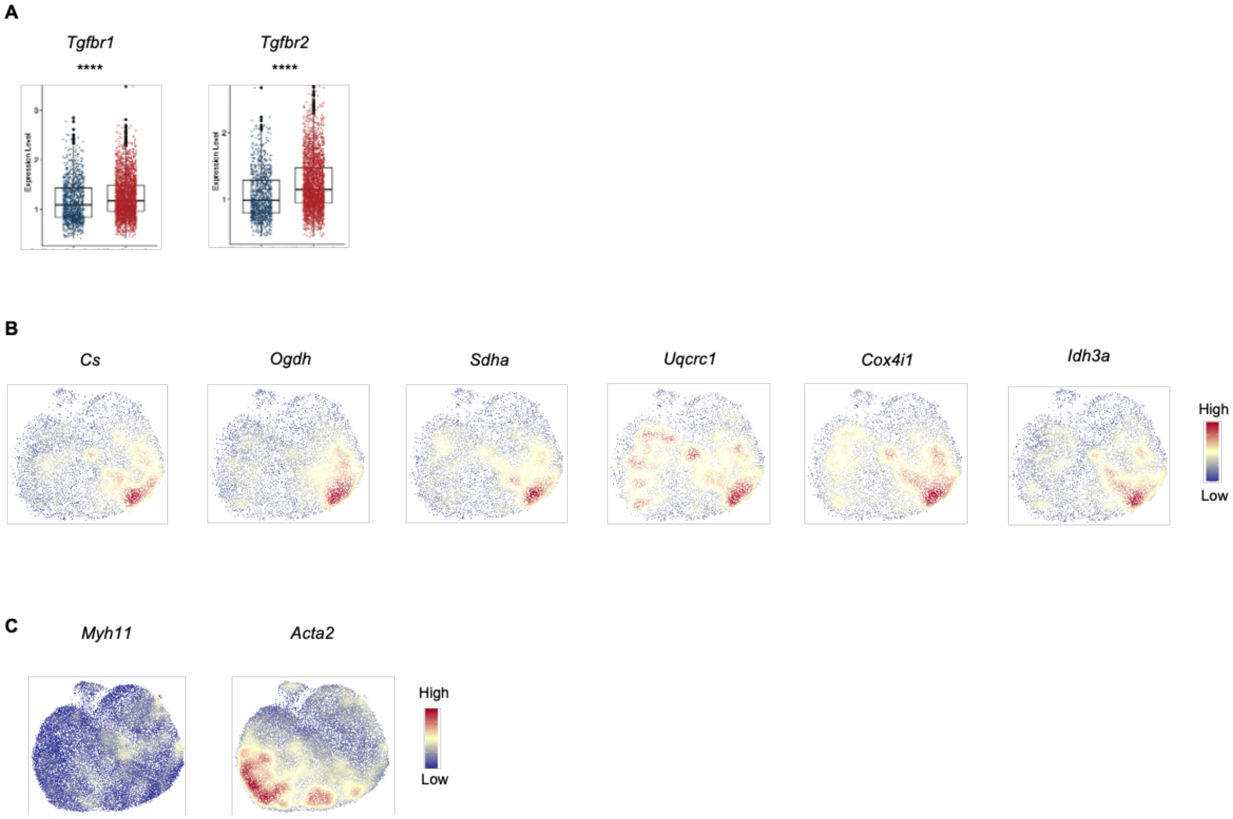


**Supplemental Figure 3. Transcriptional characterization of Fbn1-deficient mice.** (**A**) Boxplots quantifying expression of selected genes in mVIC1 comparing +/+ and *C1039G*/+ valves. Data are shown as individual data points with mean ± SEM. ****P < 0.0001, by Wilcoxon test. (**B**) UMAP showing expression of representative oxidative phosphorylation (OXPHOS) and tricarboxylic acid (TCA) cycle-related genes across VIC clusters. (**C**) UMAP showing expression of canonical myofibroblast markers across VIC clusters.

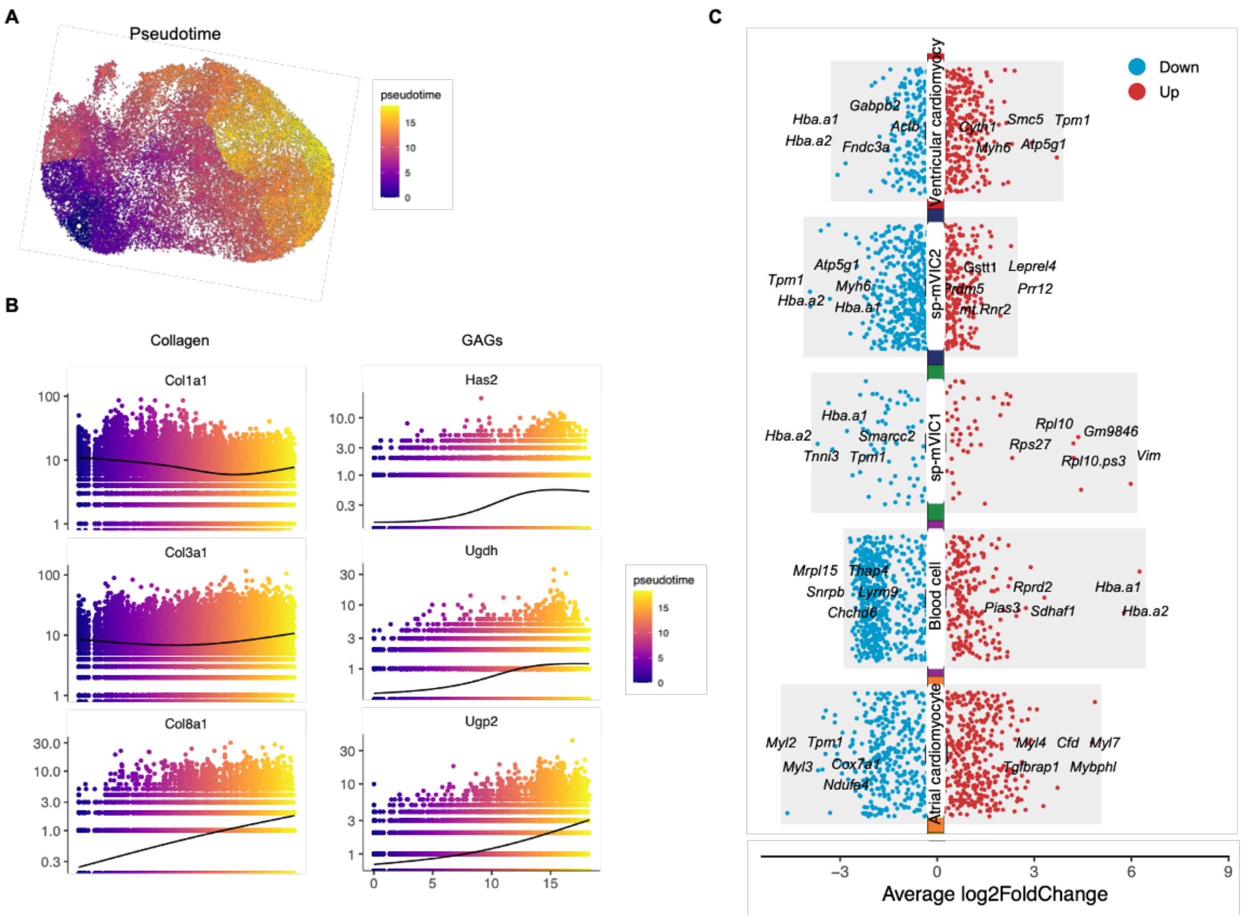


**Supplemental Figure 4. Characterization of Fbn1-deficient mice.** (**A**) UMAP of VICs overlaid with inferred pseudotime trajectories, illustrating predicted transcriptional progression across VIC states. (**B**) Scatter plots showing dynamic expression patterns of representative genes involved in collagen synthesis and GAG regulation along the pseudotime trajectory. (**C**) Scatter plots depicting DEGs across spatial transcriptomic clusters, with the top five DEGs labeled. Genes highlighted in red are upregulated, whereas those in blue are downregulated.

A

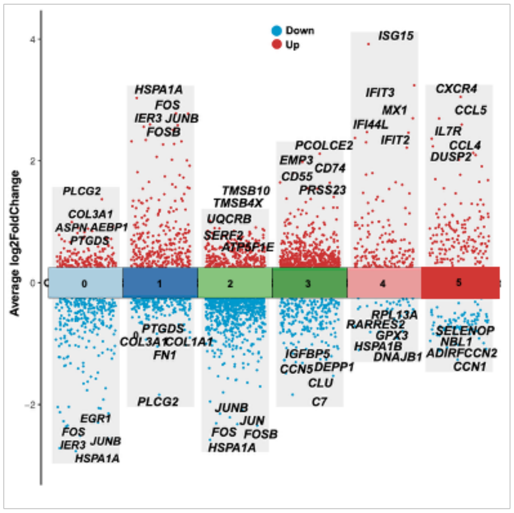
VIC
Average log2FoldChange
Down
Up
0
1
2
3
4
5
ISG15
HSPA1A
FOS
IER3 JUNB
FOSB
IFIT3
MX1
IFI44L
IFIT2
CXCR4
CCL5
IL7R
CCL4
DUSP2
PCOLCE2
EMP3
CD74
CD55
PRSS23
PLCG2
COL3A1
ASPN AEBP1
PTGDS
TMSB10
TMSB4X
UQCRB
SERF2
ATP5F1E
PTGDS
COL3A1 COL1A1
FN1
PLCG2
RPL13A
RARRES2
GPX3
HSPA1B
DNAJB1
SELENOP
NBL1
ADIRF CCN2
CCN1
IGFBP5
CCN5 DEPP1
CLU
C7
JUNB
JUN
FOS FOSB
HSPA1A
EGR1
FOS
IER3 JUNB
HSPA1A

B

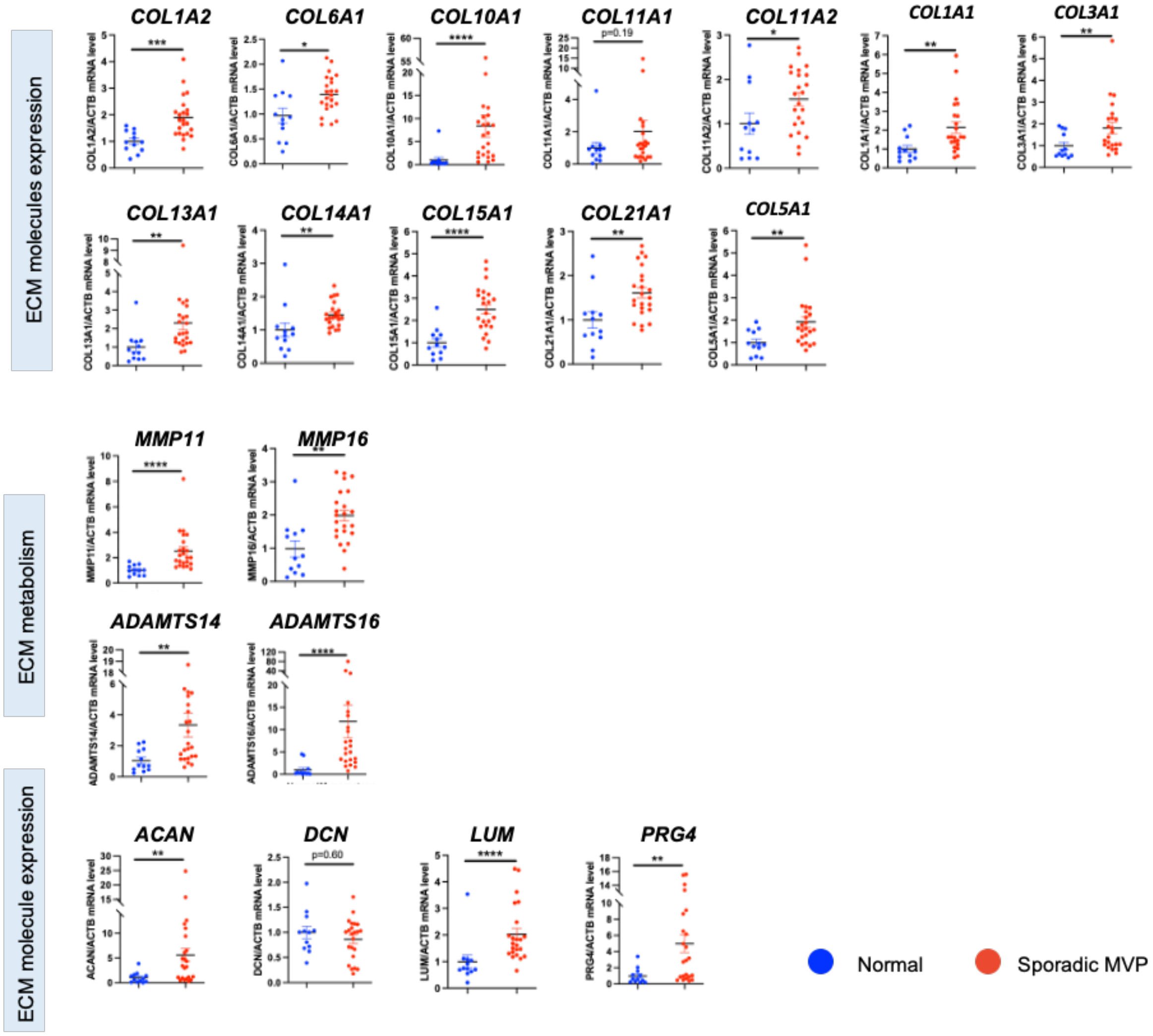
ECM molecules expression
COL1A2
COL6A1
COL10A1
COL11A1
COL11A2
COL1A1
COL3A1
COL13A1
COL14A1
COL15A1
COL21A1
COL5A1
p=0.19
ECM metabolism
MMP11
MMP16
ADAMTS14
ADAMTS16
ECM molecule expression
ACAN
DCN
LUM
PRG4
p=0.60
Normal
Sporadic MVP

**Supplemental Figure 5. Extended transcriptional characterization of sporadic MVP.** (**A**) Scatter plots depicting DEGs across VIC clusters, with the top five DEGs labeled for each cluster. Genes highlighted in red are upregulated, whereas those in blue are downregulated. (**B**) Quantitative RT-PCR analysis of gene expression in normal and sporadic MVP mitral valves. Data are presented as individual data points with mean ± SEM. Statistical significance was assessed using unpaired t test. $^{*}P < 0.05$; $^{**}P < 0.01$; $^{***}P < 0.001$; $^{****}P < 0.0001$.

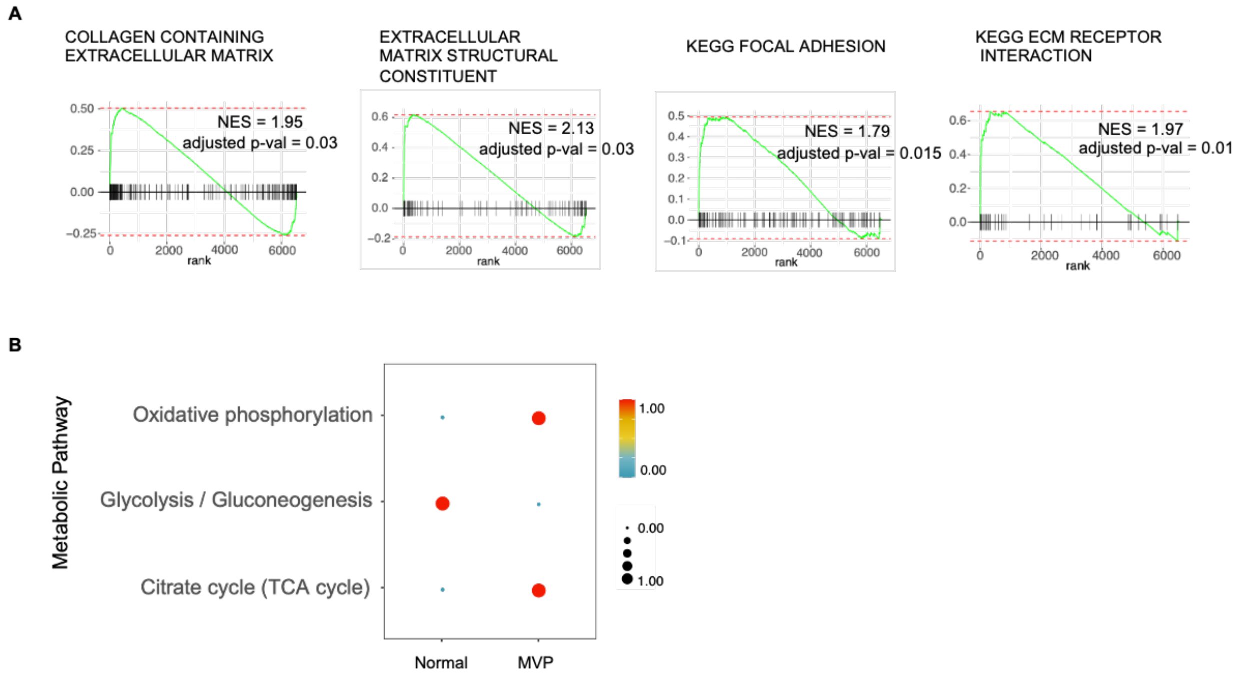


**Supplemental Figure 6. Gene set enrichment and metabolic pathway analysis of sporadic MVP.** (**A**) Gene set enrichment analysis (GSEA) performed on DEGs identified in the activated VIC cluster from sporadic MVP compared with normal mitral valves. (**B**) Comparative analysis of metabolic pathway activity in the activated VIC cluster from normal and sporadic MVP valves.

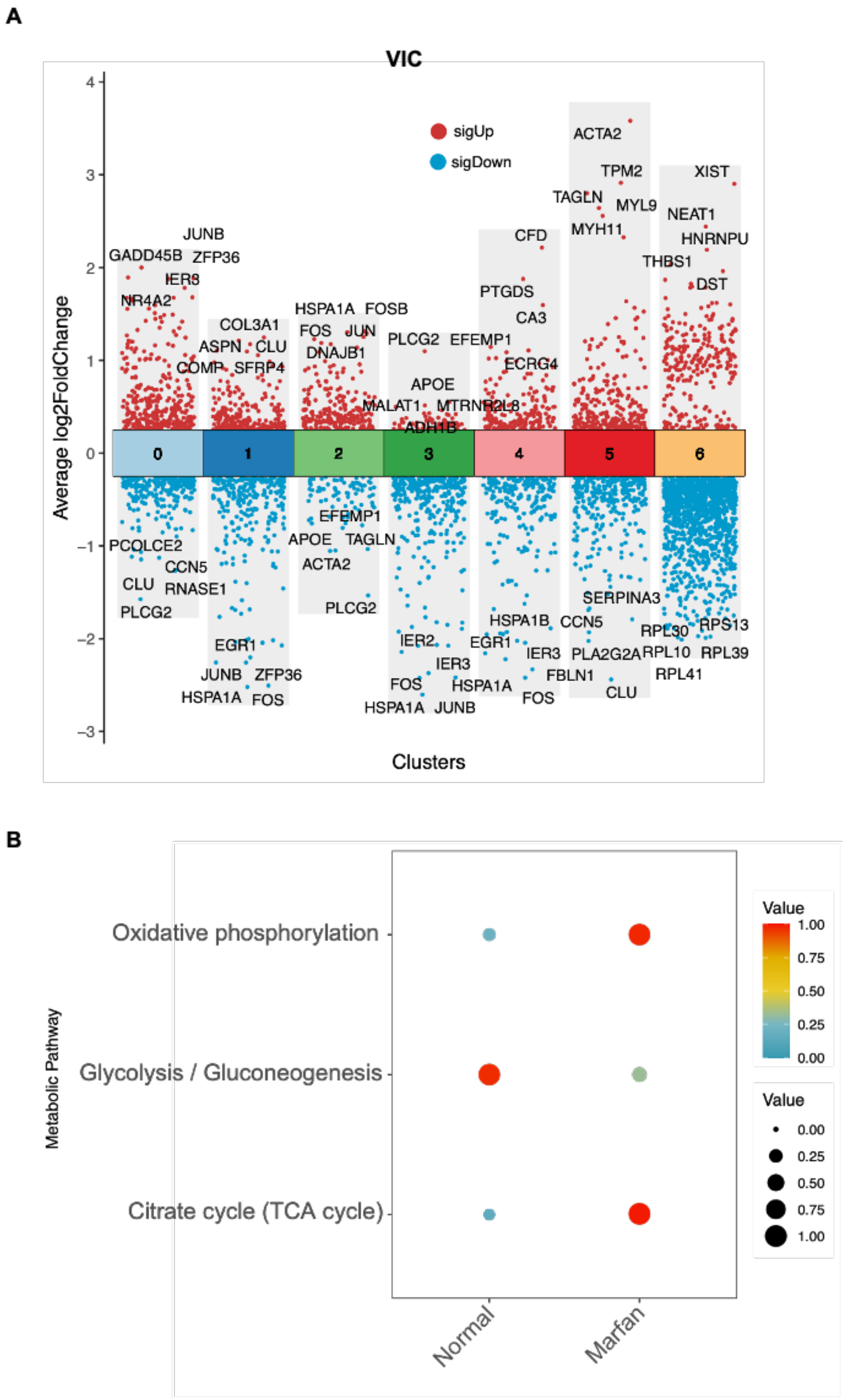


**Supplemental Figure 7. Extended transcriptional characterization of Marfan-associated MVP.** (**A**) Scatter plots depicting DEGs across VIC clusters, with the top five DEGs labeled for each cluster. Genes highlighted in red are upregulated, whereas those in blue are downregulated. (**B**) Comparative analysis of metabolic pathway activity in the activated human VICs from normal and Marfan-associated MVP valves.

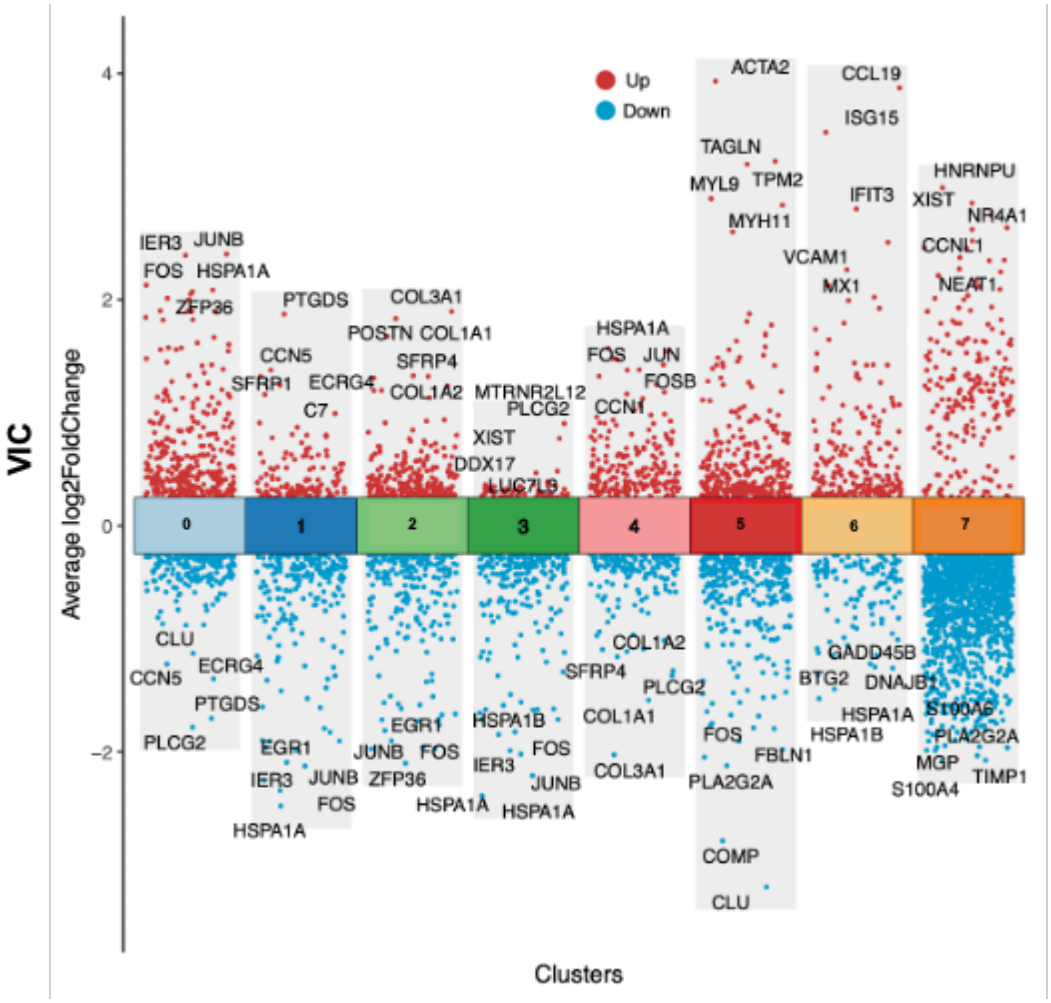


**Supplemental Figure 8. Extended transcriptional characterization of human MVP.** Scatter plots depicting DEGs across VIC clusters, with the top five DEGs labeled for each cluster. Genes highlighted in red are upregulated, whereas those in blue are downregulated.

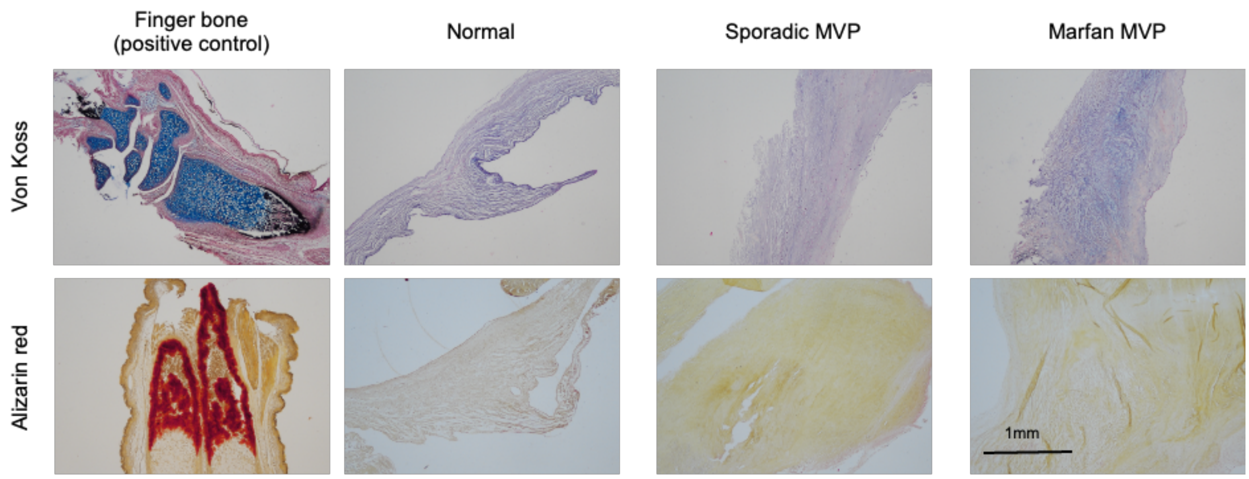


**Supplemental Figure 9. Assessment of calcium deposition in human mitral valve tissues.** Representative histological staining for calcium in human mitral valve sections, with murine finger bone included as a positive control.

**Supplemental Table 1. SenMayo senescence gene panel.**

| | | | | | |
|---|---|---|---|---|---|
| Acvr1b | Cd55 | Esm1 | Igfbp7 | Mmp10 | Scamp4 |
| Ang | Cd9 | Ets2 | Il10 | Mmp12 | Selplg |
| Angpt1 | Csf1 | Fas | Il13 | Mmp13 | Sema3f |
| Angptl4 | Csf2 | Fgf1 | Il15 | Mmp14 | Serpinb3a |
| Areg | Csf2rb | Fgf2 | Il18 | Mmp2 | Serpine1 |
| Axl | Cst10 | Fgf7 | Il1a | Mmp3 | Serpine2 |
| Bex3 | Ctnnb1 | Gdf15 | Il1b | Mmp9 | Spp1 |
| Bmp2 | Ctsb | Gem | Il2 | Nap1l4 | Spx |
| Bmp6 | Cxcl1 | Gmfg | Il6 | Nrg1 | Timp2 |
| C3 | Cxcl10 | Hgf | Il6st | Pappa | Tnf |
| Ccl1 | Cxcl12 | Hmgb1 | Il7 | Pecam1 | Tnfrsf11b |
| Ccl2 | Cxcl16 | Icam1 | Inha | Pgf | Tnfrsf1a |
| Ccl20 | Cxcl2 | Icam5 | Iqgap2 | Pigf | Tnfrsf1b |
| Ccl24 | Cxcl3 | Igf1 | Itga2 | Plat | Tubgcp2 |
| Ccl26 | Cxcr2 | Igfbp1 | Itpka | Plau | Vegfa |
| Ccl3 | Dkk1 | Igfbp2 | Jun | Plaur | Vegfc |
| Ccl4 | Edn1 | Igfbp3 | Kitl | Ptbp1 | Vgf |
| Ccl5 | Egf | Igfbp4 | Lcp1 | Ptger2 | Wnt16 |
| Ccl7 | Egfr | Igfbp5 | Mif | Ptges | Wnt2 |
| Ccl8 | Ereg | Igfbp6 | Mmp13 | Rps6ka5 | |

**Supplemental Table 2. Human mitral valve specimen used for scRNA-seq***

| Specimen | Age (yrs) | Sex | Race | Location | Disease state |
|---|---|---|---|---|---|
| **D1** | 48 | M | Caucasian | anterior | normal |
| **D2** | 48 | M | Caucasian | posterior | normal |
| **D3** | 42 | F | Caucasian | anterior | normal |
| **D4** | 42 | F | Caucasian | posterior | normal |
| **D5** | 51 | F | Caucasian | anterior | normal |
| **D6** | 51 | F | Caucasian | posterior | normal |
| **D7** | 55 | M | Caucasian | anterior | normal |
| **D8** | 55 | M | Caucasian | posterior | normal |
| **P1** | 62 | M | Hispanic | posterior | sporadic MVP |
| **P2** | 58 | M | Caucasian | posterior | sporadic MVP |
| **P3** | 65 | M | Caucasian | posterior | sporadic MVP |
| **P4** | 58 | M | Caucasian | posterior | sporadic MVP |
| **P5** | 49 | M | Caucasian | posterior | sporadic MVP |
| **P6** | 59 | M | Caucasian | posterior | sporadic MVP |
| **P7** | 70 | F | Caucasian | anterior | Marfan MVP |
| **P8** | 70 | F | Caucasian | posterior | Marfan MVP |
| **P9** | 34 | M | African American | anterior | Marfan MVP |
| **P10** | 34 | M | African American | posterior | Marfan MVP |

*Mitral valve specimen were obtained from organ donors without disease or after surgical mitral valve repair for sporadic mitral valve prolapse (MVP) or mitral valve replacement for Marfan MVP. D: donor; P: patient; M: male; F: female; yrs: years

**Supplememental Table 3. Human mitral valve specimen used for spatial RNA-seq***

| Specimen | Age (yrs) | Sex | Race | Location | Disease state |
|---|---|---|---|---|---|
| **D1** | 48 | M | Caucasian | posterior | normal |
| **P1** | 58 | M | Caucasian | posterior | Sporadic MVP |

*Mitral valve specimen were obtained from organ donors without disease or after surgical mitral valve repair for sporadic mitral valve prolapse (MVP). D: donor; P: patient; M: male; yrs: years

**Supplemental Videos Legends**

**Supplemental Video 1.** High-resolution micro-computed tomography of mouse finger bone showing positive control for calcium

**Supplemental Video 2.** High-resolution micro-computed tomography of normal mitral valve

**Supplemental Video 3.** High-resolution micro-computed tomography of Sporadic MVP mitral valve

**Supplemental Video 4.** High-resolution micro-computed tomography of Marfan mitral valve
